# Nuclear deformability facilitates apical nuclear migration in the developing zebrafish retina

**DOI:** 10.1101/2024.04.04.588091

**Authors:** Mariana Maia-Gil, Maria Gorjão, Roman Belousov, Jaime A. Espina, João Coelho, Ana P. Ramos, Elias H. Barriga, Anna Erzberger, Caren Norden

**Author notes:** Correspondence to Caren Norden. Equal contribution (Maria Gorjão, Roman Belousov).

## Abstract

Nuclear positioning is an important aspect of cell and developmental biology. One example is the apical positioning of nuclei in retinal and other neuroepithelia. Here, apical nuclear migration is crucial for correct tissue formation. Cytoskeletal mechanisms that drive nuclei to the apical side have been explored. Yet, whether also nuclear properties influence apical nuclear migration remained comparatively less understood. Lamin A/C expression levels have been shown to be directly related to nuclear deformability. Further, it was shown that many nuclei in early development, including neuroepithelial nuclei, express only low levels of Lamin A/C.

Thus, we asked whether increased expression of Lamin A in the densely packed zebrafish retinal neuroepithelium affects nuclear migration phenomena. We find that overexpressing Lamin A in retinal nuclei of single cells or in the whole tissue increased nuclear stiffness and consequently impaired apical positioning. Interestingly, also nuclei of control cells embedded in a Lamin A overexpressing environment displayed impaired apical nuclear migration. When Lamin A is overexpressed at the tissue level this further leads to a delay in mitotic entry. Thus, nuclear material properties, within cells but also in the surrounding environment, can influence nuclear and cell behavior in densely packed neuroepithelia.

Overall, this work quantitatively shows a relevance of low Lamin A/C levels in early neuroepithelial development. These findings are most likely also applicable for other developing tissues which feature nuclear and cell motion through crowded environments.

## Introduction

Nuclear positioning plays important roles in diverse contexts of successful cell and tissue development and in homeostasis^1–4^. In unicellular organisms like fission yeast as well as in tissues as pseudostratified epithelia, nuclei need to be correctly positioned before mitosis^5–10^. Further, in many migrating cells like fibroblasts or developing neurons nuclei must be moved within cells to ensure overall cell movement ^6,7,11^.

Considering this plethora of nuclear positioning phenomena and their relevance, it is unsurprising that nuclear-positioning defects can have severe organismal consequences including muscle disorders^12^ and diverse forms of central nervous system malformations^13^.

Forces to move nuclei with and within cells is generated by the cytoskeletal elements microtubules and/or actin. The exact cytoskeletal machinery involved depends on cellular and tissue context^4^. Microtubules move nuclei for example in and with diverse migrating neurons^6,14,15^. Actin-dependent mechanisms drive the rearward nuclear movement during fibroblasts polarization^11^ and move nuclei apically before division in developing neuroepithelia^8,16,17^.

In addition to the cytoskeleton, nuclear properties can influence nuclear movements with and within cells ^18^. As nuclei are the biggest and among the stiffest organelles in most eukaryotic cells, their movements can be affected by their large size and elastic properties^19–23^. *In vitro* studies of neutrophil-like cells, mouse embryonic fibroblasts and diverse cancer cell-lines showed that nuclear deformability influences nuclear movements through microchannels^19,24–28^. How much nuclei were able to deform depended on the expression levels of the intermediate filament Lamin A/C. Lamin A/C are two isoforms of a nuclear envelope protein which expression levels determine nuclear shape, size and stiffness^29^. Nuclear deformability negatively correlates with Lamin A/C expression levels^24,28,30^. When Lamin A/C levels were genetically reduced in mouse embryonic fibroblasts or breast-cancer cells, this led to increased nuclear deformability and improved cell migration in confinement^24,28,29^. Conversely, overexpression of Lamin A/C in lung and breast cancer cells made nuclei less deformable and was sufficient to delay confined cell migration^28,31^. The combination of this work was very insightful; however most studies were so far performed using *in vitro* settings. It is currently much less explored whether and how nuclear properties linked to Lamin A/C expression influence tissue development and homeostasis *in vivo*. During development, cells and their accompanying nuclei frequently need to move through confined environments as seen for example during gastrulation and in other morphogenesis frameworks^13^. Thus, it is tempting to speculate that also here nuclear properties linked to Lamin A/C expression influence such movements. This notion gets more traction when considering the fact that in many settings of early vertebrate development, naturally occurring Lamin A/C expression levels are low to non-detectable. This is true for example in the zebrafish early retinal neuroepithelium^16^, in the early mouse brain^32^ and in mouse and human embryonic stem cells^33^. Thus, that low Lamin A/C levels could facilitate nuclear and cell migration phenomena in developing and often crowded tissues that undergo diverse morphogenetic rearrangements.

To probe this notion, we here used the zebrafish retinal neuroepithelium as a model tissue. Neuroepithelia are densely packed pseudostratified tissues consisting of elongated cells that are apically and basally attached. Nuclei in these cells move during the cell cycle (Video S1): In G1 nuclei passively drift to basal positions where they oscillate during S phase. In G2 nuclei actively and directedly move apically for mitosis^5,8–10,34^. These G2 movements and subsequent apical mitosis are crucial to guarantee tissue integrity and further maturation^1,2^. It has been shown that in retinal neuroepithelia apical nuclear movement is driven by formin-dependent actin polymerization where actin filaments are not directly linked to the nuclear envelope^16^. Additionally, it was observed that only low levels of Lamin A/C are expressed in these retinal neuroepithelia^16,35^. We thus asked whether overexpression of Lamin A could influence nuclear deformability and mobility. Indeed, Lamin A overexpression leads to less deformable nuclei that show impaired apical migration in G2. This effect is seen when all nuclei overexpress Lamin A, but also when overexpression occurs in a cell autonomous manner in otherwise control tissues. Interestingly, also in the converse scenario where control nuclei expressing low levels of Lamin A are exposed to a Lamin A overexpression environment, these control nuclei move less efficiently. Furthermore, onset of mitotic rounding is delayed in a Lamin A overexpression environment. Together this means that Lamin A expression levels can influence nuclear properties during tissue development and thereby nuclear and cell behavior in cell autonomous but also non-cell autonomous manners.

## Results

### Lamin A overexpression in zebrafish retinal neuroepithelia decreases nuclear volume and influences the ability for shape changes

Since it has been shown that retinal neuroepithelial nuclei express low levels of Lamin A/C^16^, we tested if nuclear properties changed when we overexpress Lamin A using a Tg(hsp70:LMNA-mKate2) transgenic line^36^. Lamin A overexpression (OE) was induced by heat shocking embryos at 22 hours-post fertilization (hpf) at 39 °C for 30 minutes^36^.

To confirm that overexpression was successful, zebrafish heads were dissected 6 hours after heat-shock and Lamin A protein levels were analyzed by Western Blot. While Lamin A is almost absent in controls, as reported previously for immunostainings^16^, detection is much increased upon heat-shock treatment (Figure 1 A). The increase of Lamin A levels did not impact the protein levels of Lamin B, another main component of the nuclear lamina (Figure S1 A).

**Figure 1.**
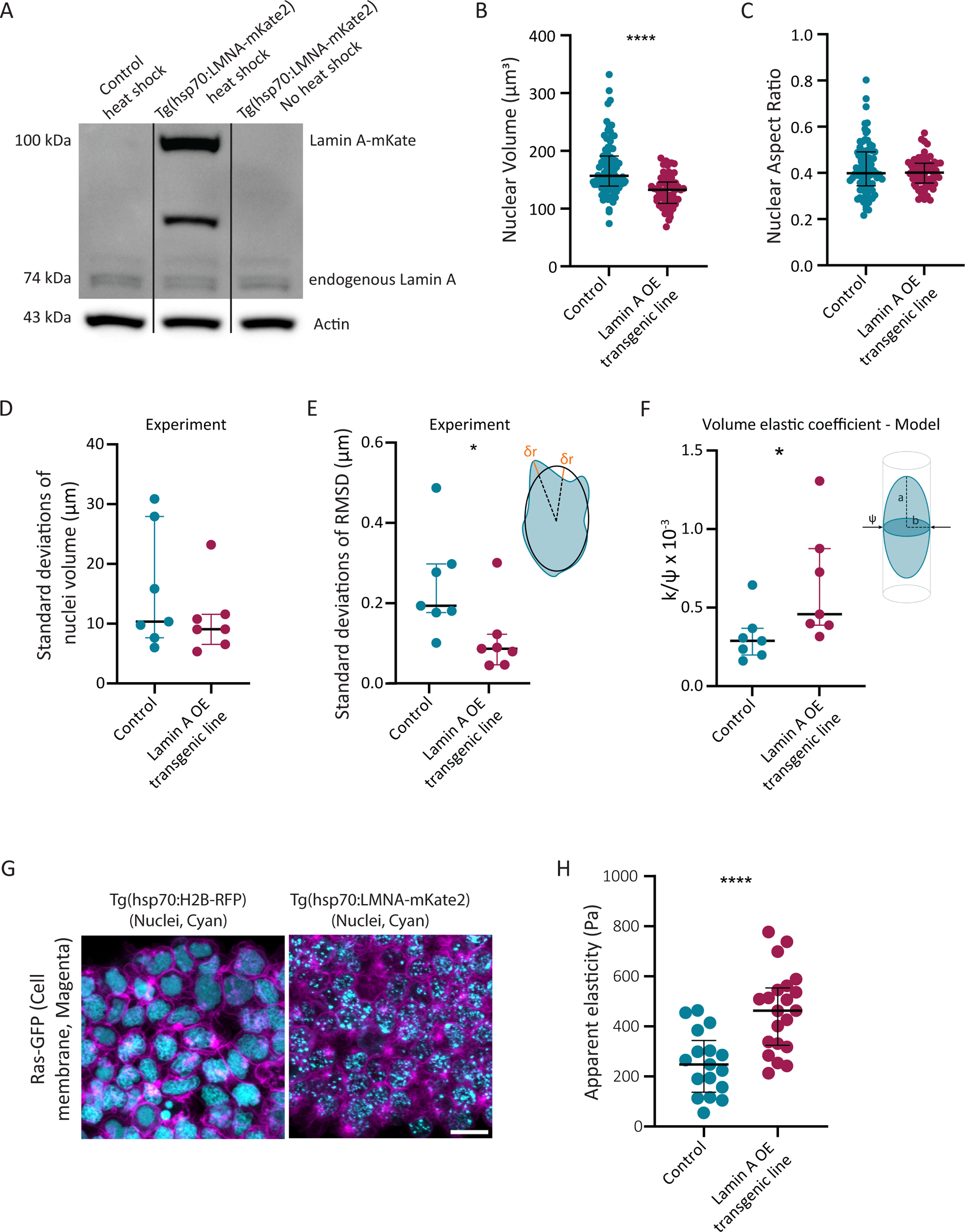
Lamin A overexpression changes nuclear and tissue properties in retinal neuroepithelia. (A) Western Blot detecting Lamin A levels in control Tg(hsp70:H2B-RFP) after heat shock (left), Lamin A OE line Tg(hsp70:LMNA-mKate2) after heat-shock (middle) and Lamin A OE line Tg(hsp70:LMNA-mKate2) without heat-shock (right). (B,C) Quantification of nuclear volume (B) and aspect ratio (C) of 3D segmented nuclei (see Figure S1B). Nuclei overexpressing Lamin A are isometrically smaller when compared to controls (P_Volume_ < 0.0001, P_Aspect Ratio_ = 0.5891, Mann-Whitney test). Variance of nuclei volume and aspect ratio is less in Lamin A OE transgenic line (Variance comparison: P_Volume_ < 0.0040, P_Aspect Ratio_ < 0.00119, Levene’s test). Error bars: Median and interquartile range. (D) Standard Deviations (SD) of the nuclei volume in S phase at consecutive time-points (P_SD_Volume_= 0.457, Mann-Whitney test), (see Figure S1D). (E) Standard Deviations (SD) of the Root Mean Squared Deviation (RMSD) of S phase nuclei at consecutive time-points (P_SD_RMSD_= 0.026, Mann-Whitney test), (see Figure S1E). Scheme or Root Mean Squared Deviation (RMSD) analysis in the right top corner. (F) Nuclei volume elastic coefficient inferred from nuclear segmentations in Figure S1C and D (P_Volume Elastic Coeficient_ = 0.011, Mann-Whitney test). Schematic of parameters used in theoretical model (see Supplementary Note) in the right top corner. (G) Retinal cells dissected and plated on a glass-bottom dish. Tg(hsp70:H2B-RFP) and Tg(hsp70:LMNA-mKate2) respectively label nuclei (cyan), RAS-GFP labels the plasma membrane (magenta). Note that control and Lamin A OE nuclei are labeled by different nuclear markers, H2B-RFP and LMNA-mkate2 respectively, which explains different patterns. Scale bar: 10 μm. (H) Atomic Force Microscopy (AFM) measurements of plated retinal cells shown in G, (P < 0.0001, t-test). Error bars: Median and interquartile range.

To probe whether Lamin A OE affects nuclear architecture and shape, we mosaically labelled nuclei with H2B to analyze nuclear volume. Nuclei were segmented in 3D and their volume assessed in control and Lamin A OE retinas (Figure S1 B). Control nuclei showed an average volume of 169.8 ± 47.83 µm^3^, whereas Lamin OE nuclei were approximately 20% smaller at an average volume of 130.4 ± 27.47 µm^3^ (Mean± SD) (Figure 1 B). In both conditions, nuclei presented ellipsoidal shapes elongated along the apicobasal (AB) cell axis and showed similar aspect ratios (Figure 1 C). Overall, nuclear volumes and aspect ratios were less variable across Lamin A overexpressing nuclei than across controls (Figure 1 B and C). To analyze whether Lamin A OE also affected nuclear deformability, we mosaically labelled nuclei with lap2b, a nuclear envelope marker, and segmented nuclear shapes over time in both control and Lamin A OE retinas (Figure S1 C). We then evaluated nuclear volume (Figure S1D) and the root mean squared deviations (RMSD, Figure 1 D) of their surface from a perfect ellipsoid (Figure S1 C). RMSD analysis gives an appreciation of the response of the whole nucleus to spontaneous perturbations and quantifies fine-scale surface effects which gives insights into overall small-scale deformations. We initially performed this analysis over S phase as this is the phase in which nuclei do not actively move along the AB axis^9^, meaning that the observed shapes during this period correspond to near-equilibrium configurations. This analysis constitutes a novel approach to quantify the fine-scale morphology of nuclei, which permits characterising nuclear deformability directly from imaging data. This expansion was necessary for the type of analysis conducted here, as so far most studies probing the effect of nuclear properties on nuclear and cell migration were done using single cell *in vitro* or more qualitatively *in vivo* analysis^22,24,26,28^. The result of this analysis revealed that most control nuclei displayed a larger amplitude of surface fluctuations than the Lamin A OE nuclei as quantified by the standard deviations calculated over each trajectory (Figure E and S1E). This means that the shape response to random forces is more resilient in Lamin A OE nuclei. Together this shows that Lamin A OE nuclear are smaller and less deformable than their control counterparts.

### Lamin A overexpression changes nuclear and tissue stiffness: theory and experiment

We next used our experimental data and RSDM analysis and developed a simple mechanical description of the confined nuclei as compressible droplets to understand a possible effect of Lamin A expression on nuclear stiffness. For this, we approximated the nuclear surface as a prolate axisymmetric spheroid with two degrees of freedom - a longer apicobasal axis of length *a* and two shorter axes of length *b*, governed by a coarse-grained effective free energy *F*(*a*, *b*) = *γA*(*a*,*b*) + *k*2[*V*(*a*,*b*)−*V*_0_]2−*ψ α*(*b*), (1) in which *V*(*a*,*b*), *A*(*a*,*b*), *α*(*b*) are, respectively, the spheroid’s volume, surface area, and the area of its largest cross-section perpendicular to the apicobasal axis (Figure 1 E, S1 C and E). The surface tension *γ* > 0 represents the free-energy cost of the droplet’s interface with the environment^37,38^. The volume elastic coefficient *κ* > 0 penalizes deviations of the nuclear volume from its preferred value *V*_0_ > 0, similar to the cellular volume elasticity commonly used in vertex or cellular-Potts and other soft matter models^39–44^. Finally, we introduced a coefficient *ψ* that is conjugate to the cross-sectional area of the nucleus to effectively account for the stress transversal to the apicobasal axis: when *ψ*=0, the equilibrium configuration of the nucleus corresponds to a perfect sphere of radius *a*=*b*=*γ*/*ΔP* given by the Young-Laplace law with the Laplace pressure *ΔP*=*δF*/*δV*=*k* [*V*(*a*,*b*)−*V*_0_], whereas the prolate spheroid *a*>*b* equilibrates for *ψ*>0. Due to the competing effects of surface tension and transversal stress, the preferred volume is expected to exceed the equilibrium volume.

To estimate the parameter space of retinal nuclei, we used the time series of their segmented 3D shapes in S phase (Figure 1E, S1 C and E). We assumed these shapes to correspond to near-equilibrium configurations, because nuclei do not actively move during this phase^9^. By least square fitting (see Supplementary Note) we determined the parameter values *γ*/*ψ*, *κ*/*ψ*, and *V*_0_ for control and Lamin A OE nuclei from measurements of their ellipsoidal axes. As expected from the experimental analysis of nuclear volumes (Figure 1 B) also theoretical analysis found smaller preferred volumes for Lamin A nuclei (885.7±182.8 µm^3^) compared to the control sample (1056 ± 360 µm^3^) (Figure S1 F). Also, the volume elastic coefficient estimated for the two nuclei populations differed almost two-fold, with κ/ψ increasing from (3.2 ± 1.6) × 10^−5^ µm^−4^ in the control sample to (6.4 ± 3.6) × 10^−5^ µm^−4^ for the Lamin A OE cells (Mean ± SD) (Figure 1 F). Further, the surface tensions also increased: from 0.26 ± 0.07 *ψ* to 0.52 ± 0.21 *ψ* in the control and Lamin A OE populations, respectively (Mean ± SD) (Figure S1 G) (for a more complete description of the model we refer the reader to the Supplementary Note).

Overall, our theoretical analysis predicts an increase in the volume elastic coefficient. This in turn suggests that overexpression of Lamin A increases the resistance of nuclei to compression, meaning they should be stiffer.

To corroborate the theoretics predictions and extract the apparent elastic moduli of retinal tissues and experimentally we used Atomic Force Microscopy (AFM) indentations. Due to the compacted disposition of nuclei in the retina which are the biggest organelle (Figure 1 G), we expected a change in nuclear properties to affect tissue-scale stiffness. For this, retinas of control embryos expressing the nuclear marker H2B and retinas of embryos overexpressing Lamin A were dissected and retinal clusters were probed with AFM *ex vivo* (Fig.1 G, Fig. S1 H-J and Material and Methods). These *ex vivo* indentations revealed that Lamin A overexpressing samples displayed a two-fold increase in their apparent elastic moduli as compared to control tissues (Fig. 1 H, Figure S1 G-I and Material and Methods).

Taken together, theoretical analysis and experimental measurements confirm that Lamin A overexpression leads to stiffer and less deformable nuclei.

### Overexpressing Lamin A specifically lengthens the G2 phase of the cell cycle

Having determined that Lamin A OE impacts nuclear properties i.e., volume and stiffness, we asked if this would have any effect on cell cycle dynamics or cell cycle dependent nuclear movements along the apicobasal axis. Cell cycle dependent movements in G1 and S phase are of non-active, stochastic nature while movements in G2 are directed to the apical surface before mitosis.

We initially excluded that Lamin A OE leads to DNA damage, which by itself could lead to changes in cell cycle kinetics, as no difference in the appearance of γ-H2AX-positive cells between control and Lamin A OE retinas was observed (Figure S2 A). To test other effects of Lamin A OE on the cell cycle, we used light sheet microscopy due to its reduced phototoxicity^45–47^. Zebrafish retinas of control and Lamin A OE embryos were imaged at 1 min intervals during the proliferative phase (24-36 hours post-fertilization (hfp)) which features only scarce neurogenic events^48^. Nuclei in both conditions were labelled mosaically using a proliferating cell nuclear antigen (PCNA) marker. This live cell cycle marker labels replication foci during S phase and allows for the unambiguous identification of all cell cycle phases^9^. We found that the overall cell cycle length (Figure 2 A) as well as G1 and S phases (Figure 2 B and C, respectively) were similar between control and Lamin A OE nuclei. However, the length of G2 increased significantly in Lamin A OE embryos (Figure 2 D). While control nuclei took on average 28.64 ± 9.10 min in G2, nuclei overexpressing Lamin A spent 43.43 ± 19.28 min in G2 (mean ± SD). However, this 14.79 min or 151% increase of G2 did not significantly lengthen the overall cell cycle of Lamin A OE cells as G2 is the shortest phase of the cell cycle (Figure 2 A-D)^9,48^. To understand if this lengthening is cell autonomous, a mosaic condition in which only single cells overexpress Lamin A upon DNA plasmid injection was introduced. Also in this context, in which single cell nuclei in an otherwise control environment overexpress Lamin A, the lengths of the overall cell cycle, S and G1 matched those of controls (Figure 2 A-D). However, also here G2 was lengthened, on average by 11.79 min or 140% compared to controls (Figure 2 D).

**Figure 2.**
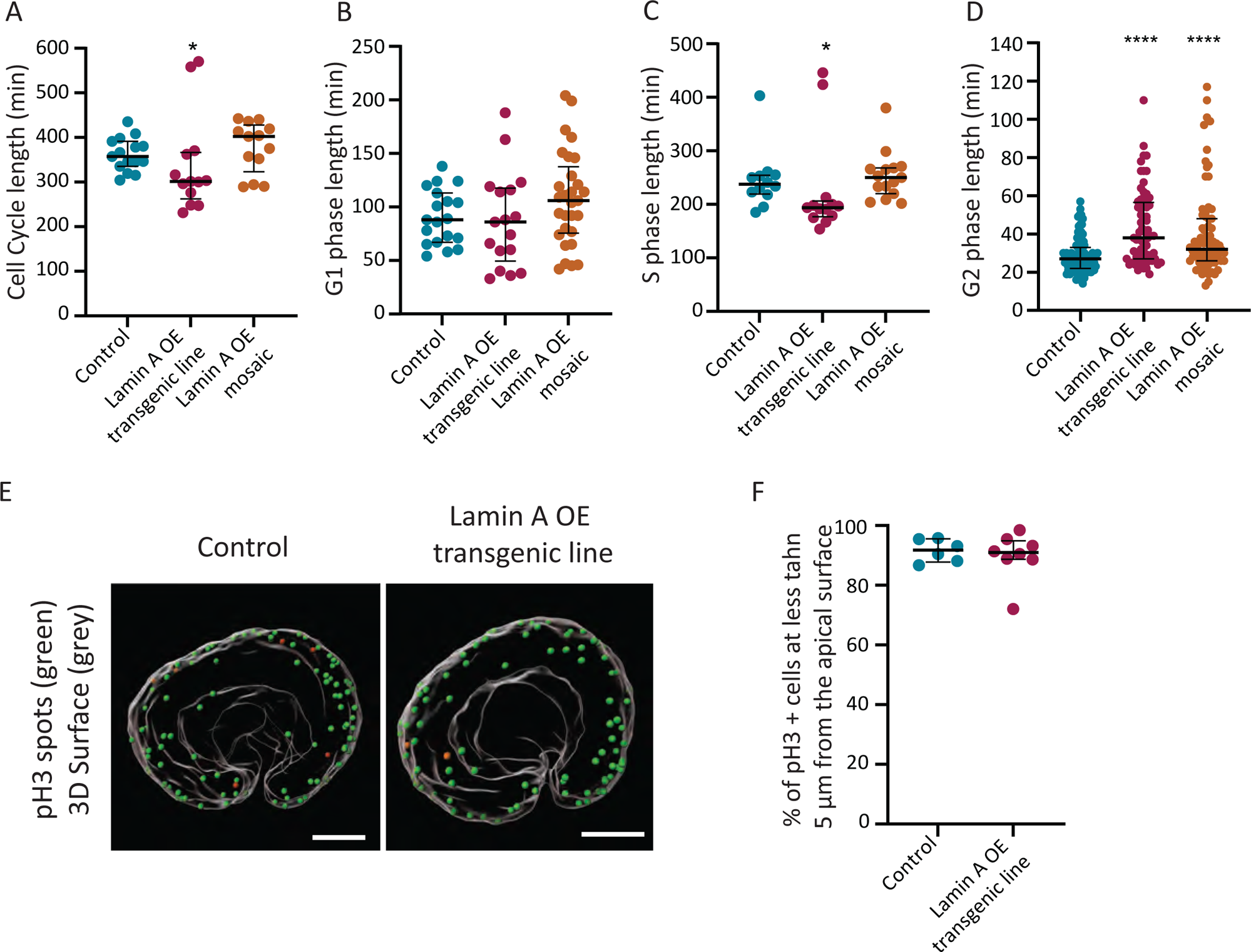
Lamin A overexpressing nuclei spend more time in G2. (A-D) Duration (in minutes) of cell cycle (A), G1 (B), S (C) and G2 (D) phases in control, Lamin A OE line and Lamin A OE mosaic retinal neuroepithelial cells. Imaged at 1 min intervals. (Cell cycle length: P_LaminA OE transgenic line_ = 0.0245, P_Lamin A OE mosaic_ = 0.3701, Mann-Whitney test and t-test, respectively; G1: P_Lamin A OE transgenic line_ = 0.8337, P_Lamin A OE mosaic_< 0.1302, t-test; S phase: P_Lamin A OE transgenic line_ = 0.0188, P_Lamin A OE mosaic_ = 0.3792, Mann-Whitney test; G2: P_Lamin A OE transgenic line_ < 0.0001, P_Lamin A OE mosaic_< 0.0001, Mann-Whitney test;). (E) pH3 spot detection of mitotic cells in the neuroepithelium of controls and Lamin A OE transgenic line. 3D retinal surface (grey). pH3 spots were classified according to their distance to the 3D surface: Green: pH3 spots within ≤ 5 μm from the apical surface; Orange: pH3 spots > 5 μm from the apical surface (right). Scale bar: 50 μm. See Figure S2B. (F) Quantification of the % of pH3 spots within ≤ 5 μm (green) from the apical surface. (P_% pH3 spots_> 0.9999, Mann-Whitney test).

Combined, our results indicate that overexpression of Lamin A specifically increases the duration of the G2 phase of the cell cycle without significantly changing the overall cell cycle length, in a cell-autonomous manner.

### Nuclei overexpressing Lamin A show slower and less efficient apical migration before mitosis

The lengthening of G2 and only G2 upon Lamin A OE was intriguing. As noted above, G2 is the only cell cycle phase in which nuclei actively move apically in a directed manner^9^. That nuclei overexpressing Lamin A nevertheless reach the apical surface to divide was shown by phospho-H3 immunostaining. This analysis showed that 90% of nuclei overexpressing Lamin A in late G2 or mitosis were within a < 5µm distance from the apical surface, compared to 92% in controls (Figure 2 E-F and Figure S2 B). Interestingly, Lamin A OE nuclei, in mosaic and transgenic line conditions, follow similar trajectories as control nuclei in S phase (Figure S3 A-D) and G1 phase (Figure S3 E-H).

To understand whether Lamin A OE could affect nuclear trajectories and thereby lead to a lengthening of G2, nuclei were followed from the moment they entered G2, marked by the disappearance of PCNA dots, until they reached the apical surface^16^ in the different conditions (Figure 3 A-F and Video S2). Nuclei overexpressing Lamin A took longer to reach the apical surface: 30.62 ± 17.46 min in the Lamin A OE transgenic line and 36.95 ± 23.38 in the Lamin A OE mosaic condition, compared to 20.32 ± 8.39 min in controls (mean ± SD) (Figure 3 G).

**Figure 3.**
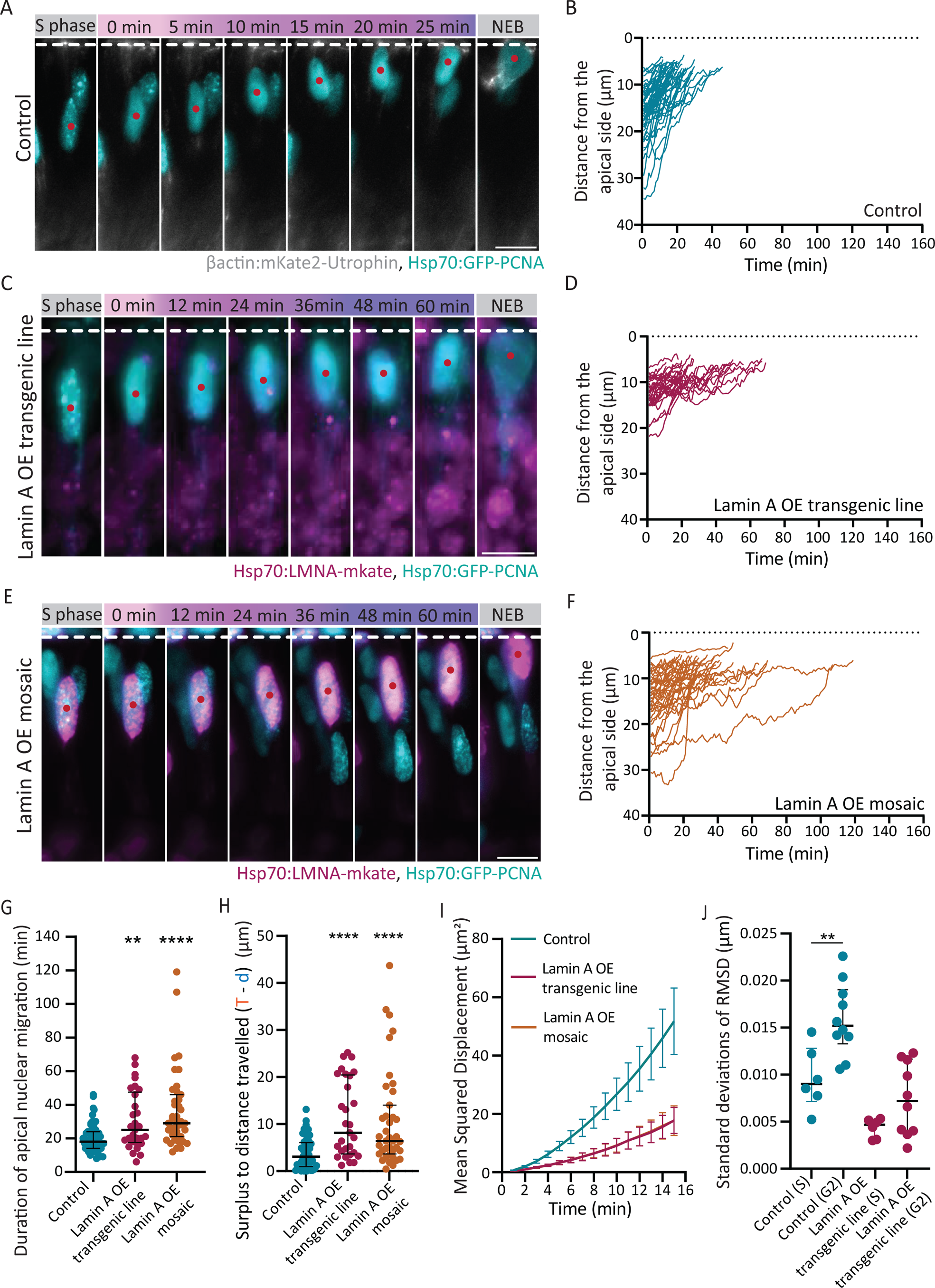
Nuclei overexpressing Lamin A are slower and less directed during apical migration. (A, C and E) Example montages of apical nuclear movements in control (A), Lamin A OE transgenic line (C) and Lamin A OE mosaic (E) retinal neuroepithelia. Red dots indicate the nucleus followed. Dashed line corresponds to the apical surface. Gradient bar from pink to purple indicates the time of apical nuclear migration, where darker colors represent longer time spans. Hsp70:PCNA-GFP labels nuclei depending on cell cycle stage (cyan), hsp70:LmnA-mKate2 labels nuclei overexpressing Lamin A (magenta). Note that in C nuclei labelled by a red dot overexpress Lamin A, although the LMNA-mkate marker is not visible in the montage. NEB: nuclear envelope breakdown. Scale bar: 10 μm. (B, D and F) Apical migration trajectories for control nuclei (B, cyan), Lamin A OE line (D, magenta), Lamin A OE Mosaic (F, orange). Dashed line corresponds to the apical surface. (G) Duration (in minutes) of apical nuclear migration, defined as the time between G2 onset and the time nuclei reach the apical side (P_Lamin A OE transgenic line_ = 0.008, P_Lamin A OE mosaic_ < 0.001, Mann-Whitney test). (H) Surplus traveled distance corresponding to the difference between the total path covered (T) and the shortest distance between starting point and apical position (d). See d and T in Figures S4B and S4C, respectively (P_Lamin A OE transgenic line_ < 0.001, P_Lamin A OE mosaic_ < 0.001, Mann-Whitney test). (I) Mean Squared Displacement (MSD) analysis in control (cyan), Lamin A OE in the transgenic line (magenta) and Lamin A in the mosaic condition (orange). MSD curves for Lamin A OE transgenic line and Lamin A OE mosaic condition mainly overlap. (J) Plot of the Standard deviations of the RMSD trajectories during G2 phase (see Figure S4E) compared to S phase (see Figure 1E and S1E) (P_Control_ = 0.003, P_Lamin A OE line_ = 0.147, Mann-Whitney test). Error bars: Median with interquartile range.

We next asked whether this delay could be explained by Lamin A OE nuclei having to move longer distances before reaching apical positions. However, we found that, in the mosaic condition, nuclei overexpressing Lamin A started apical migration at similar distances from the apical side to what was seen in controls, 14.64 ± 5.96 µm compared to 16.04 ± 5.80 µm. In the transgenic line, nuclei overexpressing Lamin A started apical migration, on average, slightly closer to the apical side, at 12.52 ± 3.60 µm (Mean ± SD) (Figure S4 A). Thus, the shortest distance between the starting and division positions ((d) in Figure S4 B) was not increased upon Lamin A overexpression. We further tested whether nuclei overexpressing Lamin A move less directedly. Interestingly, when we analyzed the surplus of traveled distance, i.e., the difference between the total path covered ((T), Figure S4 C) and the shortest distance between starting and division positions ((d)), Figure S4 B), we observed that the surplus distance for control nuclei is on average 3.79 ± 3.29 µm but for nuclei overexpressing Lamin A expands to 10.75 ± 10.43 µm in the transgenic Lamin A OE line, and 11.02 ± 8.18 µm in the Lamin A OE mosaic condition (Mean ± SD) (Figure 3 H). This increase indicates that nuclei overexpressing Lamin A exhibit less directed movements and move back and forth more frequently during apical migration. Mean square displacement (MSD) analysis showed a similar trend (Figure 3 I). Furthermore, nuclei overexpressing Lamin A displayed, on average, lower instantaneous velocities (0.19 ± 0.66 µm/min in the transgenic line and 0.22 ± 0.68 µm/min for the mosaic condition) compared to controls (0.48 ± 0.72 µm/min) (Mean ± SD) (Figure S4 D).

This means that Lamin A OE changes nuclear behavior when all nuclei overexpress Lamin A (transgenic Lamin A OE line) but also when overexpression occurs in single cells in an otherwise control environment (Lamin A OE mosaic condition). In both cases, Lamin A OE nuclei are slower and move apically in a less efficient manner than control nuclei. These data also show that nuclear properties can influence a specific stage of the cell cycle here seen in G2 lengthening.

### Nuclei overexpressing Lamin A deform less than controls during apical nuclear migration

We next asked whether the less efficient and slower apical migration of Lamin A OE nuclei could be linked to the fact that these nuclei are stiffer (Figure 1 E-H), and therefore undergo less deformations during apical migration. Control and Lamin A OE nuclei were once more mosaically labelled by the nuclear envelope marker lap2b (Figure S1 C). Nuclei were segmented in 3D and nuclear envelope shape changes were followed during apical migration (Figure S4 E). We then calculated how the RMSD diverged from the perfect ellipsoid in G2, during apical nuclear migration, as done for S phase in Figure 1E (Figure 3 J). Control nuclei in G2 undergo a larger amplitude of deformations than in S phase, as seen by the standard deviations (SD) of the displacement trajectories (Figure 3 J and Figure S4 E), and as suggested previously^16^. Interestingly, this does not apply for the amplitude of nuclear deformations in nuclei overexpressing Lamin A. Here, the already less deformable nuclei in S phase (Figure 1 E) showed a very similar RMSD amplitude in G2 (Figures 3 J, and S4 E). This indicates that, unlike controls, Lamin A OE nuclei, already stiffer in S (Figure 1 E), do not show increased deformation during G2, the period of active migration. Thus, the less efficient migration of Lamin OE nuclei to the apical side is likely linked to their increased stiffness and, consequently, lower deformability.

### Control nuclei show slower nuclear apical migration when transplanted into a Lamin A overexpressing environment

We have thus far shown that nuclei overexpressing Lamin A are less deformable and migrate apically with reduced efficiency in the transgenic and mosaic conditions.

However, the stiffness of the surrounding microenvironment can also affect nuclear and cell migration^49^. Additionally, nuclei can sense confinement in their microenvironment and adapt their behavior^50,51^. We therefore asked whether altering nuclear stiffness in the environment using the Lamin A OE line could influence nuclear apical migration of otherwise control nuclei. To this end, we performed transplantation experiments in which donor cells from control zebrafish blastulas were transplanted into control or Lamin A OE recipients (Figure 4 A). The respective nuclei were tracked as done in control, Lamin A mosaic and transgenic line conditions (Figures 4B-4E and Video S3). We observed that the time between the onset of G2 and reaching the apical side was 21.64 ± 8.54 min for control nuclei in a control environment (very similar to controls in Figure 3 G). However, this time significantly increased to 42.29 ± 26.53 min for control nuclei that moved in a Lamin A OE environment (Mean ± SD) (Figure 4 F). Thus, overexpressing Lamin A in the environment was sufficient to prolong apical nuclear migration of transplanted control nuclei on average by 20 min or 190%.

**Figure 4.**
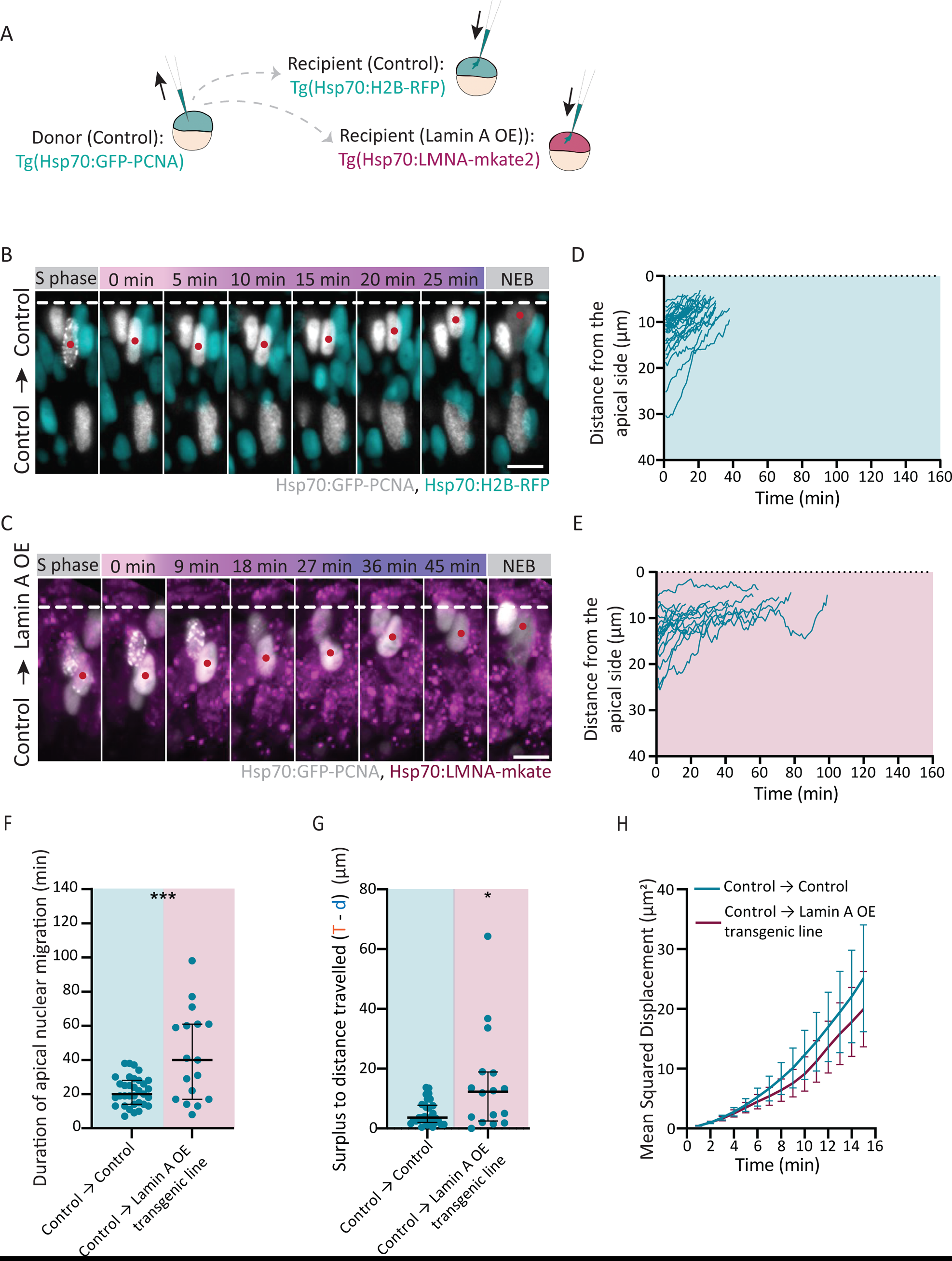
Lamin A overexpression in the environment influences apical nuclear migration of control nuclei. (A) Transplantation scheme: cells from control Tg(b-actin:PCNA-GFP) blastulas were transplanted into control Tg(hsp70:H2B-RFP) or Lamin A OE Tg(hsp70:LMNA-mKate2) blastula. (B and C) Example montage of apical nuclear migration of control cells transplanted into control (B) or Lamin A OE transgenic line (C) blastula. Red dot indicates the nucleus followed. Dashed line corresponds to the apical surface. Gradient bar from pink to purple indicates the time of apical nuclear migration, where darker colors represent longer time spans. Hsp70:PCNA-GFP labels transplanted nuclei (grey), hsp70:H2B-RFP (cyan) and hsp70:LmnA-mKate2 (magenta) show surrounding nuclei. NEB: nuclear envelop breakdown. Scale bar: 10 μm. (D and E) Apical nuclear migration trajectories for control cells transplanted into controls (D), and control cells transplanted into the Lamin A OE transgenic line (E). Dashed line corresponds to the apical surface. (F) Quantification of the duration of apical nuclear migration (min), defined as the time between G2 onset and the moment nuclei reach the apical side (P_Duration of Apical Migration_= 0.003, Mann-Whitney test). (G) Surplus travelled distance corresponding to the difference between the total path covered (T) and the shortest distance between starting point and apical position (d). See Figures S5B and S5C. (P_Duration of Apical Migration_= 0.025, Mann-Whitney test). Error bars: Median with interquartile range. (H) Mean Squared Displacement (MSD) analysis of control cells transplanted into controls (cyan) and control cells transplanted into Lamin A OE transgenic line (magenta). Error bars: Standard Deviation. Imaged at 1min time intervals.

Nuclei started apical migration at similar distances from the apical side when transplanted into control or Lamin A OE, environments (Figure. S5 A). Further, the distance between the starting and division positions (d) were similar (Figure S5 B). However, the surplus of travelled distance (T (Figure S5 C) – d (Figure S5 B)) of control nuclei in a Lamin A OE environment was on average 15.15 ± 17.09 µm, which is 3 times more than that observed for control nuclei moving in a control environment (5,08 ± 3.94 µm on average, Figure 4 G). In line with these observations, the nuclear instantaneous velocity of control nuclei was reduced in a Lamin A OE environment (0.18 ± 0.68 µm/min) when compared to a control environment (0.31 ± 0.62 µm/min) (Mean ± SD) (Figure S5 D). These results were corroborated by the respective MSD analyses (Figure 4 H). Thus, apical movements of control nuclei are less efficient when they occur in an environment in which the surrounding nuclei are stiffer due to overexpression of Lamin A.

### Lamin A overexpression in the tissue environment delays nuclear envelope breakdown and mitotic rounding

In addition to the prolonged apical nuclear migration of control nuclei in a Lamin A OE environment, we noted that these nuclei spend longer times at the apical surface before undergoing nuclear envelope breakdown (NEB) (Figure 5 A-B and Video S4) a phenomenon that further lengthened G2. While control nuclei moving in a Lamin A OE environment took on average 42.29 ± 26.53 min to reach the apical surface (Figure 5 F), G2 length was 70.94 ± 37.62 min (Mean ± SD) (Figure S5 E). This delay between reaching the apical side and displaying NEB (marking entry into mitosis and the end of the G2 phase) was also observed in the transgenic Lamin A OE line, albeit not as pronounced as in the transplantation scenario, but not in the mosaic condition (Figure 5 E). This indicated that the delay between apical arrival and NEB is related to overexpression of Lamin A in a non-cell autonomous manner and most likely linked to an increase in nuclear stiffness of the surrounding environment.

**Figure 5.**
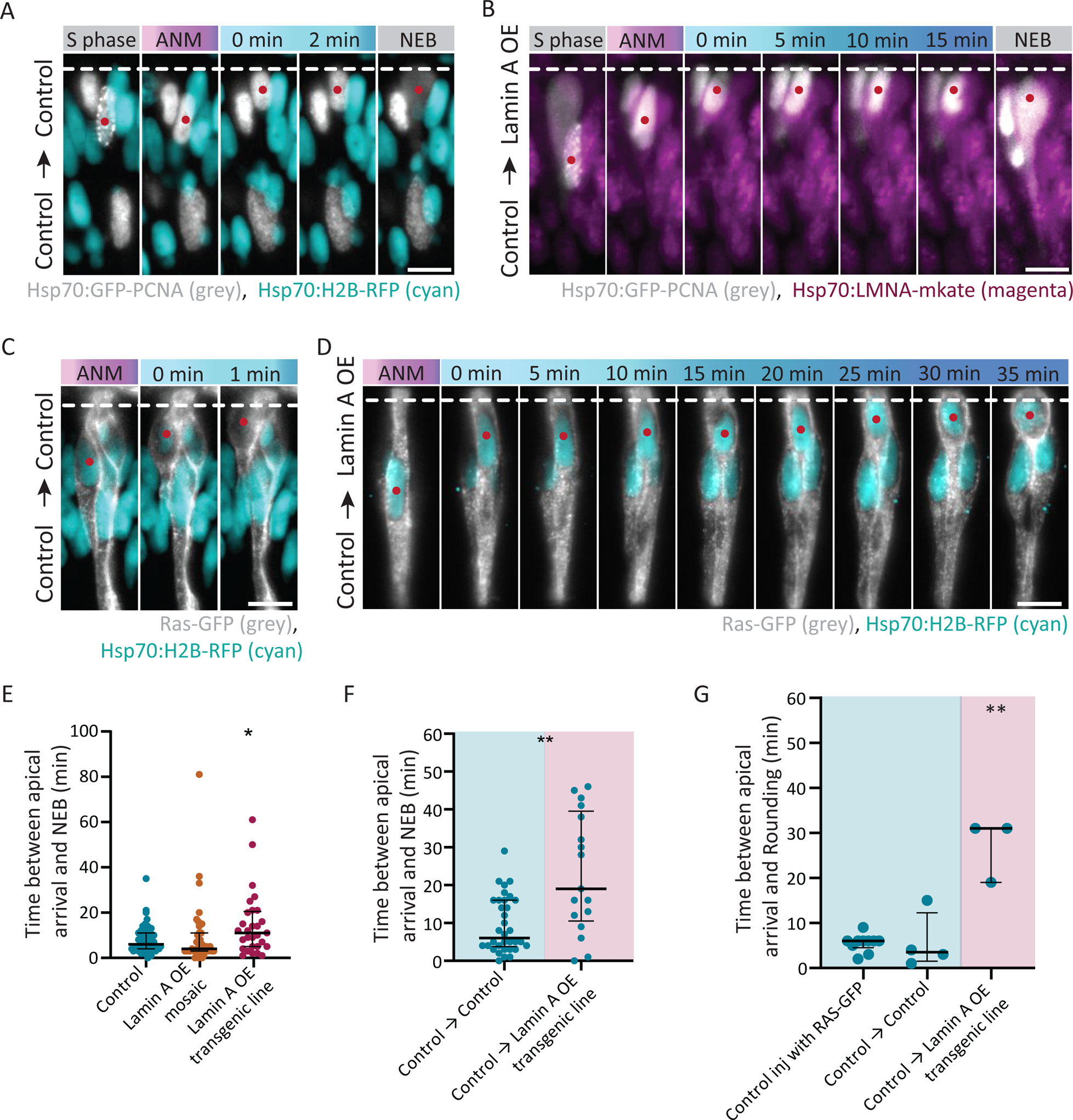
Lamin A overexpression in the environment influences timing of NEB and mitotic rounding. (A-B) Example montage of nuclear envelope breakdown (NEB) of control cells transplanted into control (A) or Lamin A OE line (B). Red dot indicates the nucleus followed. Nuclei reach the apical side at time-point = 0min. Blue bar corresponds to the time nuclei spend at the apical surface before undergoing NEB, where darker colors correspond to longer time periods. Dashed line corresponds to the apical surface. Hsp70:PCNA-GFP labels transplanted nuclei (grey), hsp70:H2B-RFP (cyan) and hsp70:LMNA-mKate2 (magenta) label surrounding nuclei. NEB: nuclear envelope breakdown. ANM: apical nuclear migration. Scale bar: 10 μm. (C-D) Example montage of mitotic rounding of control cells transplanted into control (C) or Lamin A OE line (D). Red dot indicates the nucleus followed once it reached the apical side. Dashed line corresponds to the apical surface. Hsp70:H2B-RFP (cyan) and RAS-GFP (grey) labels transplanted nuclei and plasma membrane, respectively. hsp70:H2B-RFP (cyan) label surrounding nuclei in control and the hsp70:LMNA-mKate2 label surrounding nuclei in Lamin A OE is not visible in the montage. Scale bar: 10 μm. (E-F) Quantification (in minutes) of the time between reaching the apical surface and nuclear envelope breakdown (NEB) in Control, Lamin A OE mosaic and transgenic line (E) and of control cells transplanted into control or Lamin A OE environment (F) (P_Lamin A OE mosaic_ = 0,2386, P_Lamin A OE transgenic line_ = 0,0282, P_Control → Lamin A OE line_ = 0,0027, Mann-Whitney test). (G) Quantification of the time cells take between reaching apical surface and onset of mitotic rounding. (In comparison with Control → Lamin A OE line: P _Control_ = 0,007, P _Control → Control_ = 0,0571, Mann-Whitney test). Error bars: Median with interquartile range.

Concurrently with NEB, cells need to generate space for mitotic cell rounding^52^. We therefore asked if a stiffer Lamin A OE environment could influence mitotic rounding in a non-cell autonomous manner as seen for the onset of NEB. To assess this, control cells were followed in control or Lamin A OE environments (Figures 5C-5D and Video S5). We observed that control cells in a control environment started to round immediately after their nuclei had reached the apical surface (5.75 ± 6.29 min). In contrast, control cells surrounded by cells which nuclei overexpress Lamin A took about 27 ± 6.93 min between apical arrival and the onset of rounding (Mean ± SD) (Figure 5 C, D and G). This shows that changing tissue properties by increasing Lamin A expression not only perturbs apical nuclear migration but also delays mitotic onset.

## Discussion

This quantitative study using experiment and theory tested how nuclear properties, linked to expression of the nuclear envelope protein Lamin A, influence nuclear migration and mitotic entry *in vivo*.

We find that in the densely packed zebrafish retinal neuroepithelium overexpression of Lamin A changes nuclear properties, making nuclei less deformable and increasing overall tissue stiffness. One major consequence of this increase of nuclear stiffness are changes of kinetics and overall efficiency of apical nuclear migration before mitosis. This effect was observed not only when all cells in the tissue overexpressed Lamin A, but also when single cells overexpressed Lamin A in an otherwise control tissue. This means that stiffening of a nucleus in a single cell hampers migration behavior of this nucleus. In addition, also when control nuclei move apically in a Lamin A overexpressing environment apical nuclear migration is impeded meaning that changes in the stiffness in the surrounding cells can directly affect movements of otherwise control and thereby deformable nuclei. Thus, overexpression of Lamin A influences apical nuclear migration in a cell- and in a non-cell autonomous fashion.

While a link between Lamin A/C expression and cell migration in confined environments has been made previously, most studies have so far been performed *in vitro*, often using cell migration assays through microchannels^24,25,28,31,51^. We here expand on these findings by unveiling a direct connection between Lamin A expression and nuclear movements and mitotic entry in the tissue context during embryonic development. The phenomenon depicted here, apical nuclear migration before mitosis, is relevant in all pseudostratified epithelia studied so far, as it guarantees tissue integrity for further retinal development^1,2^. Thus, our results strongly suggest that there can be a relevance of low Lamin A/C expression in developing tissues.

We are aware that Lamin A overexpression could also affect nuclear properties besides their stiffness, such as chromatin arrangements as previously described in other systems^53,54^. However, we do neither observe overall changes in cell cycle dynamics nor DNA damage upon Lamin A OE. The fact that the only cell cycle phase that lengthens upon Lamin A expression is G2, which is the only cell cycle phase in which nuclei move directly towards the apical surface, suggests that particularly the apical migration phenomenon is affected. This means that nuclear properties can directly influence cell cycle events, in this case G2 length. This lengthening of G2 in the crowded neuroepithelia upon Lamin A OE is likely independent of signaling cascades as it also occurs for control nuclei moving in Lamin A overexpressing environments. Despite the less efficient migration, however, in all scenarios nuclei reach the apical side and divide there. As apical division is a prerequisite for continued and successful development of pseudostratified epithelia^1,2^ it is likely that apical nuclear migration must be performed in a very robust manner and persist despite obstacles and perturbations. Thus, the possibility to lengthen G2 allows also the less deformable Lamin A OE nuclei to reach the apical surface before mitotic entry. This indicates that and the end (apical mitosis) justifies the means (longer apical migration and longer G2) as previously suggested in a different scenario linked to cytoskeletal machineries^16^. It will be interesting to investigate if nuclear properties can be associated with cell cycle dynamics in other morphogenesis settings and how this would influence development and organogenesis.

The very low expression levels of Lamin A in controls seem to allow nuclei to undergo extreme deformations (Video S6^35^). During the cell cycle, deformations occur most pronouncedly during apical nuclear migration in G2^16^ (Figure 4). Our data shows that these deformations can aid efficient apical nuclear migration in a densely packed tissue like the retinal neuroepithelium since the ability of nuclei to deform in the surrounding environment to reach apical locations makes migration more efficient. Nuclear deformability at the tissue level can also influence apical movements as seen by the experiment that even the in principle deformable control nuclei cannot move apically efficiently when exposed to an overall stiffer environment. This suggests that also surrounding nuclei must be deformable for efficient apical migration probably to create space in the densely packed tissue. The experiments that show delayed mitotic entry in a stiffer environment of Lamin A OE expressing nuclei argue in the same direction. Overall, this shows that *in vivo* nuclear material properties can influence morphogenetic events.

A consequence of the aforementioned considerations is that the low Lamin A (and C) expression seen in diverse early developmental settings including the Drosophila wing disc pouch, mouse and human embryonic stem cells and other tissues^16,33,55^ most likely are of relevance for dynamic morphogenesis events in which cells and nuclei need to move through densely packed tissues. It will be exiting to explore this idea across development in different epithelial and non-epithelial tissues and in different developmental frameworks to test whether common parameters exist and whether and where nuclear deformability is of general importance.

However, the observed scarce expression of Lamin A/C in early development changes when development proceeds and Lamin A/C gets upregulated for example during mouse brain formation^32^ or upon mouse and human embryonic stem cell differentiation^30,56^. This increase of Lamin A/C seems to be important as reduced Lamin A/C or loss of function mutations in differentiated tissues have been linked to disease scenarios for example in the cardiac and skeletal muscle, peripheral nerve and bones^57,58^. This indicates that nuclear deformability in early development can aid successful and timely morphogenesis and tissue formation. However, once nuclei and cells reach their final position it might be advantageous to make nuclei more rigid, and thereby protecting the genomic content^57^. To explore when Lamin A/C expression levels change and how these changes are regulated during tissue development, homeostasis and in disease will be important routes for future studies.

## Material and Methods

### Zebrafish husbandry

Wild-type zebrafish (Danio rerio; AB and Tupfel long-fin (TL) strains) and transgenic lines were kept and bred at 28 °C. Embryos were raised at 28.5°C in E3 medium. Staging of larvae was performed in hours post-fertilization (hfp) according to Kimmel et al. 1995^59^. Medium was replaced daily from 8 hpf and supplemented with 0.2 mM 1-phenyl-2-thiourea (10107703, Acros Organics) to inhibit melanogenesis. Zebrafish larvae were studied between 24 and 36hpf, a time window during which only scarce neurogenesis occurs in the retina and the majority of the cells are proliferating progenitors. Anesthesia was performed by supplementing the E3 medium with 0.04% tricaine methanesulfonate (MS-222, 1004671, Pharmaq) before live imaging and larvae dissections. All animal work was performed in accordance with the European Union directive 2010/63/EU and with the Portuguese Decree Law n° 113/2013.

### Zebrafish transgenic lines used

To visualise nuclei, the Tg(hsp70:H2B-RFP) and the Tg(β-actin:PCNA-GFP) transgenic lines were used. The Tg(β-actin:PCNA-GFP) transgenic line was further used to analyze nuclear cell cycle dynamics as previously described. To induce Lamin A OE at a tissue-wide level we used the Tg(hsp70:LMNA-mKate2) zebrafish transgenic line^36^. The Tg(β-actin:Ras-GFP) transgenic line was used to visualize the plasma membrane.

### DNA and RNA injections of zebrafish embryos

To visualize zebrafish retinal nuclei in a mosaic manner, DNA constructs were injected at one-cell stage of control embryos. The injection mixture was prepared in MiliQ water at a concentration of 50 ng of DNA per μl and 0.5-1 nl of DNA was injected per embryo. A full list of the constructs used can be found in the table S1 below. 1 nl of Ras-GFP mRNA was injected at 50 ng per μl of MiliQ water at one-cell stage embryos to label plasma membrane (see Figure 1 H).

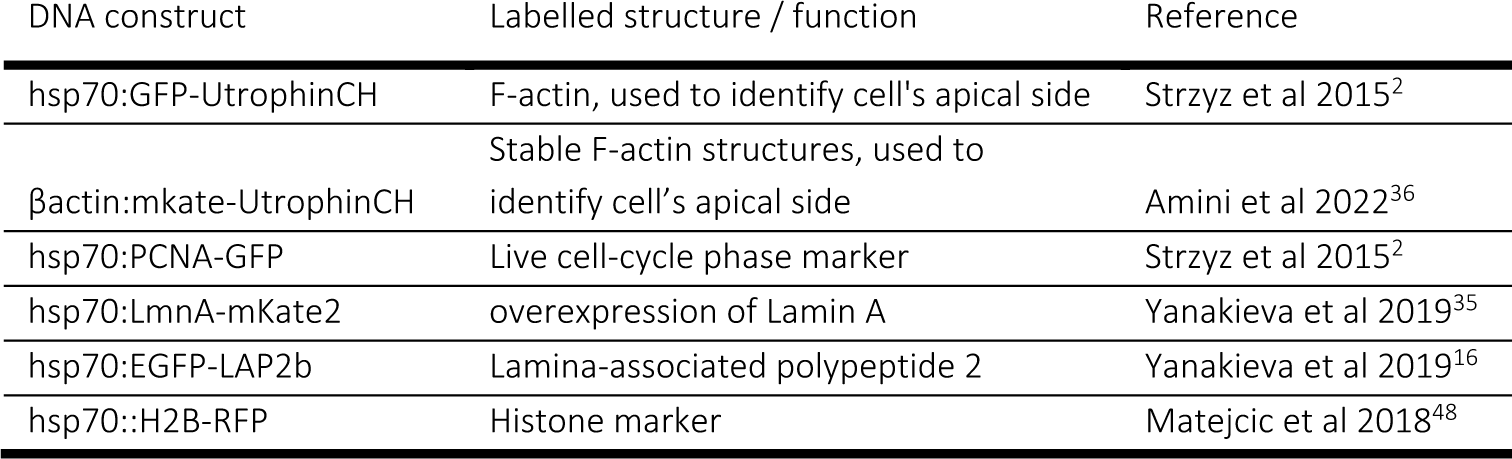

### Blastomere transplantations

To transplant blastomeres, donor (Tg(β-actin:PCNA-GFP)) and recipient (Tg(hsp70:H2B-RFP) or Tg(hsp70:LMNA-mKate2)) embryos were placed in glass-bottom dishes and dechorionated at the 256-cell stage with 2mg/ml Pronase (P8811, Sigma-Aldrich) during 5min. Embryos were rinsed 3 times with Danieus buffer (58 mM NaCl, 0.7 mM KCl, 0.4 mM MgSo4.7H2O, 0.6 mM Ca(NO3)2 and 5mM HEPES). Following dechorionation, embryos were transferred with a glass-pipette to a petri-dish, coated with 1% agarose containing 6×25 wells. Transplantations were performed in E3 medium under an Olympus SZX10 stereo microscope using a micromanipulator and a glass-needle with 0.1 mm diameter. Around 15-25 blastomere cells were transplanted per embryo during the mid-blastula stage. After transplantation, embryos were kept for 1-2 hours at 32°C. Transplanted embryos were then transferred to a 1% agarose-coated petri dish with fresh E3 medium, supplemented with 0.2 mM 1-phenyl-2-thiourea (10107703, Acros Organics) to inhibit melanogenesis, and were kept at 28.5°C over night.

### Heat-shock treatment and screening

Induction of hsp70-driven promoters was performed by incubating zebrafish larvae between 22 and 23 hpf in a water bath at 39°C for 30 min. Larvae were then kept at 28.5°C with fresh E3 medium supplemented with 0.2 mM 1-phenyl-2-thiourea (10107703, Acros Organics). Around 3-4 hours after heat-shock, zebrafish larvae were screened for the activation of the hsp70-driven promoters under a stereo microscope (Olympus SZX16) with a fluorescence lamp (Olympus U-HGLGPS). Only fluorescent positive larvae were selected to continue the experiments.

### Western-blot

#### Dissection and storage of zebrafish heads

Before dissection, zebrafish larvae at 28-30 hpf were manually dechorionated and anesthetized by supplementing the E3 medium with 0.04% tricaine methanesulfonate (MS-222, 1004671, Pharmaq). Anesthetized larvae were then placed on a glass-bottom dish with pre-cold PBS 1x. 25 heads were manually dissected with forceps (size: 55) and transferred to a 1.5 ml microtube containing minimal liquid. After dissection, samples were immediately snap-frozen on dry ice. Three repeats were done per condition, alternating between conditions in order to have comparable developmental times. Samples were stored at −80 °C before further processing.

#### Immunoblotting

Heads were lysed by adding lysis buffer [RIPA supplemented with protease inhibitors (A32953, Pierce), dithiothreitol (MB03101, NZYTech) at 50 mM and sodium orthovanadate (S6508, Merck) at 10 mM] and using an electric pestle, in 10 second bursts. After incubation on ice for 20 min, samples were heated at 95°C for 5 min and centrifuged at 18000 RCF for 10 min, 4°C. Supernatants were collected and mixed with protein loading buffer (928-40004, LI-COR Biosciences) before being loaded on pre-cast NuPAGE 4-12% Bis-Tris Protein gels (NP0322BOX, ThermoFisher Scientific). The gels were run in buffer containing 50 mM Tris base, 50 mM MOPS, 0.1% SDS and 0.1 mM EDTA (M00138, GenScript) at 100V for 200 min in a Mini Gel Tank system (ThermoFisher Scientific). The molecular weight marker used was Chameleon® Duo Pre-stained Protein Ladder (928-60000, LI-COR Biosciences).

Transfer of proteins from gels to nitrocellulose membranes (926-31092, LI-COR Biosciences) was performed using the Mini Gel Tank system, according to the recommendations of the manufacturer. Transfer was done at 20 V for 1 hour buffer containing 25 mM Tris base and 25 mM Bicine (M00139, GenScript) supplemented with 10% absolute ethanol (20821.330, VWR Chemicals).

Immunoblots were blocked for 15 min with PBS (137 mM NaCl, 2.7 mM KCl, 8.1 mM Na2HPO4 * 2H2O, 1.8 mM KH2PO4) containing 10% non-fat milk (Molico, Nestlé) and 0.05% Tween 20, followed by overnight incubation with primary antibodies, at 4 °C with slight agitation. Membranes were then washed thrice with PBS containing 0.05% Tween 20 and incubated with secondary antibodies. Following three washes with PBS containing 0.05% Tween 20 and two washes with PBS, membranes were developed using WesternBright Sirius HRP substrate (K-12043-D10, advansta), according to the manufacturer’s instructions.

#### Antibodies and antisera

For Western blot analyses, the primary antibodies used were an anti-Lamin A/C (NBP1-85550, NovusBio) at 1:1000, an anti-Lamin B1 (ab16048, abcam) at 1:2000, and an anti-actin (A2066, Sigma-Aldrich) at 1:2000. The secondary antibody used was an anti-rabbit-HRP conjugated (711-035-152, Jackson Immuno Research) at 1:10000.

Dilutions of anti-Lamin A/C and Lamin B1 antibodies were performed in buffer containing 50 mM Tris-HCl pH 7.5, 150 mM NaCl, 0.1% Triton X-100 and 2% BSA, while the anti-actin and the secondary antibodies were diluted in PSB containing 5% non-fat milk and 0.05% Tween-20.

#### Whole-mount immunostaining of zebrafish retinas

For whole-mount immunostainings, zebrafish larvae were dechorionated manually and fixed at 28-30 hpf with 4% PFA (043368-9M, Thermo Fisher Scientific) in PBS. Fixation was done overnight at 4°C. Larvae were then rinsed with PBS 1x and washed 5 times with PBS-triton 0.8% (28817295, VWR) for 10 min with slight agitation. Permeabilization was done using trypsin-EDTA (sc-391060, Santa Cruz Biotechnology) on ice for 10 min. Larvae were rinsed twice with PBS-triton 0,8% (28817295, VWR) and kept for 30 min in PBS-triton 0.8% (28817295, VWR) on ice. Blocking was performed in 10% goat serum (16210064, Gibco) in PBS-triton 0.8% (28817295, VWR) for 3 hours at room-temperature. Incubation with the primary antibody was done using 1% goat serum (16210064, Gibco) in PBS-triton 0,8% (28817295, VWR) for 3 days at 4 °C, with slight agitation. The following primary antibodies were used: 1:500 Anti-phospho-Histone H3 (Ser10) (Cat No. 631257, Sigma-Aldrich); 1:200 Histone H2A.XS139ph (phosphor-Ser139) (Cat No. GTX127342, RRDI: AB_2833105, GeneTex). After 5 washes of 30 min with PBS-triton 0.8% (28817295, VWR), larvae were incubated with the secondary antibody using 1% goat serum (16210064, Gibco) in PBS-triton 0.8% (28817295, VWR) for 2 days at 4 °C, with slight agitation. The following secondary antibodies were used: 1:500 Alexa Fluor anti-rabbit (RRID: AB_2535792, Thermo Fisher Scientific); and 1:1000 DAPI (MBD0015, Sigma-Aldrich).

### Atomic Force Microscopy

#### Dissection of retinal pseudostratified epithelium

Before retinal dissection for atomic force microscopy, zebrafish larvae at 26-30 hpf were manually dechorionated and anesthetized by supplementing the E3 medium with 0.04% tricaine methanesulfonate (MS-222, 1004671, Pharmaq). Anesthetized larvae were then placed on a glass-bottom dish with chilled PBS 1x where zebrafish heads were manually dissected. Next, the two eyes were separated from each head. To assure that we only measure the stiffness of cells from the retinal pseudostratified epithelium (PSE), the retinal pigmented epithelium (RPE) and the lens were removed from all eyes. To visualize the RPE, these larvae were not supplemented with 1-phenyl-2-thiourea (PTU), allowing for the development of melanocytes. Dissected retinal cells were transferred to a 1.5ml microtube and kept on ice. This process was repeated for 10-14 retinas per condition. After dissection, retinal cells from both conditions were platted on a glass-bottom dish, previously coated with Poly-D-Lysine (A3890401, Gibco), and kept at 28.5°C for 30min. Once cells attached to the Poly-D-Lysine, a Leibovitz’s L-15 Medium (11415049, Gibco) supplemented with 20 mM HEPES (10397023, Fisher Bioreagents) and 2% FBS (10270106, Gibco) was added to the petri-dish.

### *In vivo* Atomic Force Microscopy (AFM) measurements

All AFM measurements were done by using a FLEX-ANA (Nanosurf) automated AFM device. The AFM head was mounted in a Leica DMI 6000 inverted microscope fitted with a x–y-motorized stage that allowed to image cells by using a 10x/0,30 Leica dry objective while acquiring AFM data. Cantilevers coated with a ∼2.25 µm diameter colloidal spheres were used (CP-qp-SCONT-Au-A, NanoAndMore GmbH). This tip size ensured that the indentations capture the mechanical properties at the cellular and not the tissue level. Cantilevers were mounted on the AFM device and their spring constants were calculated using the thermal noise method^60^. Only cantilevers with spring constants between 0.01 and 0.03 N/m were selected. Extracted retinas were mounted as we previously described and 25 indentations were performed per cell in a ROI of 50×50 μm. The following modulation parameters were used: maximum indentation force, 5-7 nN; approach speed, 5 μm/s; retraction speed was 50 μm/s; sample rate, 2400 Hertz. AFM curves were selected for analysis as described below in AFM data analysis.

### AFM data analysis and image treatment

All AFM data was analysed with the software AtomicJ (v 2.3.1). Here, raw data of force– distance curves were fitted to a Hertz model for a spherical indenter,

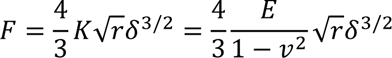

with applied force F, Young’s modulus E, Poisson’s ratio v, indenter radius r, indentation depth δ, and apparent elastic moduli K= E/(1– nu^2), referred as “stiffness” in the text and as “apparent elasticity (Pa)” in the y-axis of each chart. The quality of curves was addressed by their shape (example curve in Supplementary Figure 1b^61^). For the selected curves AtomicJ was used to detect the contact point and the elastic moduli from this point to 2 mm indentation depth was extracted in each indentation as previously described^62,63^. Then the median of each grid made on a single cluster of retinal cells was calculated and processed for further statistical analyses.

### Microscope Image acquisition

Zebrafish retinas were imaged during the proliferative phase, between 26 and 36hpf.

### Confocal scan of whole-mount zebrafish retinas

Fixed samples were imaged in a Leica Stellaris 5 upright confocal microscope using a HC PL APO 40x/1,1 Water CORR CS2 objective (#506425, Leica). The microscope was operated by the LAS X (v4.5.0.25531) software. Samples were mounted in 1% agarose in glass bottom dishes (MatTek). Acquired z-stacks had the thickness of the entire retina and step-size of 1μm.

### Time-lapse imaging using LSFM

Live imaging of single nuclei in 26 to 36 hpf zebrafish retina were performed in Zeiss Z.1 Light Sheet microscope operated with ZEN 2014 SP1 software (v9.2.8.60) (black edition). 26 hpf larvae were mounted in a 0.6% low-melting agarose (prepared in filtered E3) column and the sample chamber was filled with E3 supplemented with 0.2 mM 1-phenyl-2-thiourea (10107703, Acros Organics) and 0.04% tricaine methanesulfonate (MS-222, 1004671, Pharmaq) as previously described^45^. Entire retinas were imaged in a single-view at 1 min time intervals for 8-10 hours. Images were acquired at 0.5-1μm steps using dual-sided illumination by 10×/0.2 objectives (Carl Zeiss Microscopy). Time-lapse images were taken using the Plan-Apochromat 40×/1.0-W detection objective (Carl Zeiss Microscopy) and the two PCO.Edge 4.2 sCMOS cameras. Both retinas were image per zebrafish larvae using the multiview option.

### Image processing and analysis

Simple image processing was performed before data analysis. Using Zen (ZEN 2014 SP1 software (v9.2.8.54 and v9.3.1.393) (black edition)) or Fiji, images were averaged, cropped and drift was corrected. Image analysis was done using IMARIS (v 10.0.0) and Fiji, and results were analyzed and plotted using Microsoft Excel, GraphPad Prism 9 and Python. Statistical analysis was performed in GraphPad Prism 9.

### Quantification of apical divisions

DAPI staining of 28-30 hpf retinas was used to create a 3D retina surface using the surface tool in IMARIS. Next, the spot detection tool was used to identify nuclei positive for Ph3 immunostaining, as described above, to score mitotic nuclei. To quantify the percentage of Ph3 positive nuclei that divide apically, the spots were filtered according to the shortest distance to the created surface and classified whether they were within 5μm distance (green spots) or further (orange spots) from the apical side using IMARIS (v 10.0.0).

### Nuclear segmentation, volume and shape

Nuclei of 28-30 hpf zebrafish retinas, labelled with H2B-GFP upon injection of the hsp70:H2B-GFP plasmid at one-cell stage embryos, were segmented using the surface tool in IMARIS (v 10.1.0). 3D segmentation was achieved by fitting an ellipsoid to the fluorescently labelled nuclei. The length of the long semi axis, and of the short semi-axis, as well as the respective aspect ratios, were extracted from these 3D segmentations. Nuclear volume was obtained by calculating the number of voxels occupied by each segmented nucleus.

### Nuclear deformations during S and G2 cell cycle phases

Tg(hsp70:H2B-RFP) and Tg(hsp70:LMNA-mKate2) zebrafish were mosaically labelled with the nuclear envelope marker hsp70:EGFP-LAP2b DNA using injection at one-cell stage. Zebrafish retinas were imaged at 26-36 hpf using lightsheet microscopy as previously described. Nuclei were considered in S phase around 40-60 min before they reached the apical surface. G2 nuclei were analyzed 20 min before reaching the apical surface. Nuclear 3D segmentations were then executed using the LimeSeg plugin in imageJ (https://imagej.net/plugins/limeseg). Following, the Root Mean Squared Deviation (RMSD) of the 3D segmented nucleus was analyzed and compared to a perfect ellipsoid. The RMSD over time indicates how the nuclear shape changes in stochastic movement (S phase) and during apical nuclear migration (G2 phase). The Standard Deviations (SD) shows the amplitude of the nuclear shape changes, as a proxy for nuclear deformability.

### Analysis of nuclear dynamics during the cell cycle

Nuclei were labelled with PCNA, a live cell cycle marker that labels the replication foci, to distinguish the different phases of the cell cycle^9^. PCNA positive nuclei express replication foci during S phase. Once PCNA foci disappear the G2 phase starts and lasts until nuclear envelope breakdown (NEB). After apical mitosis, nuclei are in G1 until the reappearance of the PCNA replication foci. To quantify apical nuclear migration in the different conditions, nuclei were analyzed from the moment that G2 starts, marked by the disappearance of the PCNA foci, until when nuclei had reached the apical surface of the cell (labelled by actin marker Utrophin).

### Sample drift correction for the apical side

The cell’s apical side was considered as reference point to which nuclear positions were compared. Utrophin, that labels F-actin, was used to identify the apical surface. In the transplantation experiments, the apical surface was identified by the fact that all nuclei express H2B. Each cell crop was drift-corrected for the apical side of the cell using the Manual drift correction plugin in Fiji (https://imagej.net/plugins/manual-drift-correction).

### Maximum intensity projection, rotation and nuclei tracking

As nuclei move mainly along the y axis, and only marginally in x and z, a maximum intensity projection (MIP) of the z-stack was applied to each cell crop and rotated to align the cell axis with the y axis using Zen (ZEN 2014 SP1 software (v9.2.8.54 and v9.3.1.393) (black edition)) and/or Fiji. Therefore, each nuclear movement was analyzed in 1D, using the MtrackJ plugin in Fiji (https://imagej.net/plugins/mtrackj). For this, the center of each nucleus was manually selected in the different time-points and used as a proxy for nuclear position.

### Kinetics of apical nuclear migration

Duration and position of nuclear movements were extracted from the MtrackJ analysis and calculated in Excel (Microsoft) and Wolfram Mathematica^64^. Trajectories’ mean squared displacement (MSD) and directionality ratio were calculated from the first 18 min of apical nuclear migration or S phase and were analyzed in Excel (Microsoft) using the open-source computer program DiPer^65^.

### Statistical Analysis

All statistical tests were performed using GraphPad 9. Two-tailed tests were used and 95% confidence intervals were considered. Data was first tested for normality using the D’Agostino & Pearson and/or the Shapiro-Wilk test. Then, when data followed a Gaussian distribution, groups were compared with an unpaired t-test and, for non-parametric data, with a Mann-Whitney test. All p-values are indicated in the respective figure legend.

## Acknowledgments

We thank the Cell Biology of Tissue Morphogenesis laboratory, E. Gomes and I. Yanakieva for fruitful project and manuscript discussion. J. Gouhier and T. Ferreira are thanked for experimental help. We are grateful to the Fish Facility and the Imaging Facility of the Instituto Gulbenkian de Ciência for experimental support. We thank T. Paixao in the Advanced Data Analysis Unit of the Instituto Gulbenkian de Ciência for advice with analysis.

M. Maia-Gil was a member of the Integrative Biology and Biomedicine PhD programme. C.N. was supported by the Fundação Calouste Gulbenkian-IGC, by the European Research Council (ERC) under the European Union’s Horizon 2020 research and innovation program (No 819046) and by the Fundação para a Ciência e a Tecnologia Investigator grant (CEECIND/03268/2018). E.H Barriga was supported by an ERC Starting Grant (No 950254); EMBO Installation Grant, Project No. 4765; La Caixa Junior Leader Incoming, No. 94978 and the Fundação Calouste Gulbenkian-IGC. A.P.R. is supported by European Union’s H2020 research and innovation program under the Marie Skłodowska-Curie Actions (101038054). R.B. and A.E. acknowledge funding from the EMBL.

The authors declare no competing financial interests.

## Author contribution

C. Norden and M. Maia-Gil conceptualized work plan and methodology. M. Maia-Gil performed the majority of the experiments and analysis. M. Gorjão performed the experiments regarding the 3D nuclear segmentation analysis. A. Erzberger and R. Belousov conceptualized and performed the theoretical analysis. E.H. Barriga and J.A. Espina conceptualized and performed the atomic force microscopy experiments. J. Coelho performed the Western Blot experiments. Ana P. Ramos helped with the experiments and analysis. C. Norden and M. Maia-Gil wrote the manuscript with the help of all the other authors.

---

**Figure S1.**
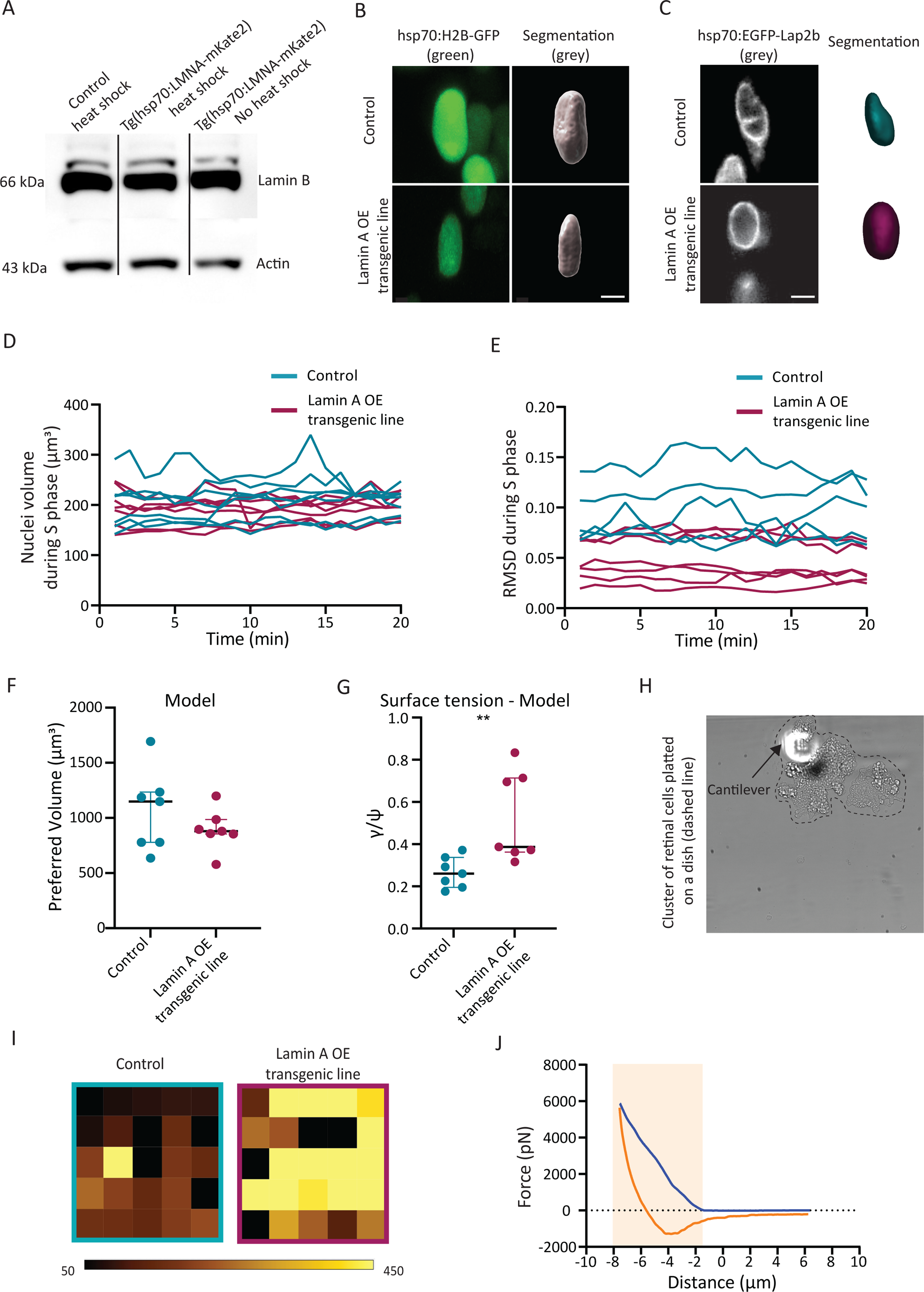
Lamin A overexpression changes tissue and nuclear properties in retinal neuroepithelia. (A) Western Blot detecting Lamin B1 levels in control Tg(hsp70:H2B-RFP) after heat shock (left), Lamin A OE line Tg(hsp70:LMNA-mKate2) after heat-shock (middle) and Lamin A OE line Tg(hsp70:LMNA-mKate2) without heat-shock (right). (B) Example of 3D segmentation of nuclei mosaically labelled with H2B-GFP in Tg(hsp70:H2B-RFP) and Tg(hsp70:LMNA-mKate2) retinas. Scale bar: 4 μm. (C) Example of 3D segmented nuclei in S phase mosaically labelled with lap2b-GFP in control Tg(hsp70:H2B-RFP) and Lamin A OE line Tg(hsp70:LMNA-mKate2). Scale bar: 5 μm. (D) Nuclei volume at consecutive time-points during S phase. 20-min trajectories showing nuclear shape changes in controls (cyan) and upon Lamin A OE (magenta). Values were used to calculate the standard deviation in Figure 1D. (E) Root Mean Squared Deviations (RMSD) of nuclear shapes from a perfect ellipsoid at consecutive time-points during S phase. 20-min trajectories showing nuclear shape changes of nuclei in controls (cyan) and upon Lamin A OE (magenta). Values were used to calculate the standard deviation in Figure 1E. (F,G) Nuclear preferred volume (F) and surface tension (G) inferred from nuclear segmentations in Figure S1C. (F: P_preferred Volume_ = 0.620; G: P_Surface Tension_= 0.004, Mann-Whitney test), (See Supplementary Note). (H) Image of the AFM cantilever position relative to the dissected retinal cells platted on a dish. (I) Heat map of control (cyan) and Lamin A transgenic line (magenta) retinal cells, showing measurements acquired in an 5×5 grid with 50 μm x/y space resolution; this grid depicts the spread of data in the tissues. All measurements used a cantilever carrying a 2.25 μm diameter bead as a tip. Each data point in the chart presented in the main figures represents a cluster of dissected retinal cells platted on a dish from which the median resulting from our 5×5 grids was calculated. (J) Representative example of a force-distance curve obtained using cantilevers coated with 2.25 μm beads. Approach curves are shown in blue and withdraw curves in orange. The indentation from the contact point is marked as light orange rectangle.

**Figure S2.**
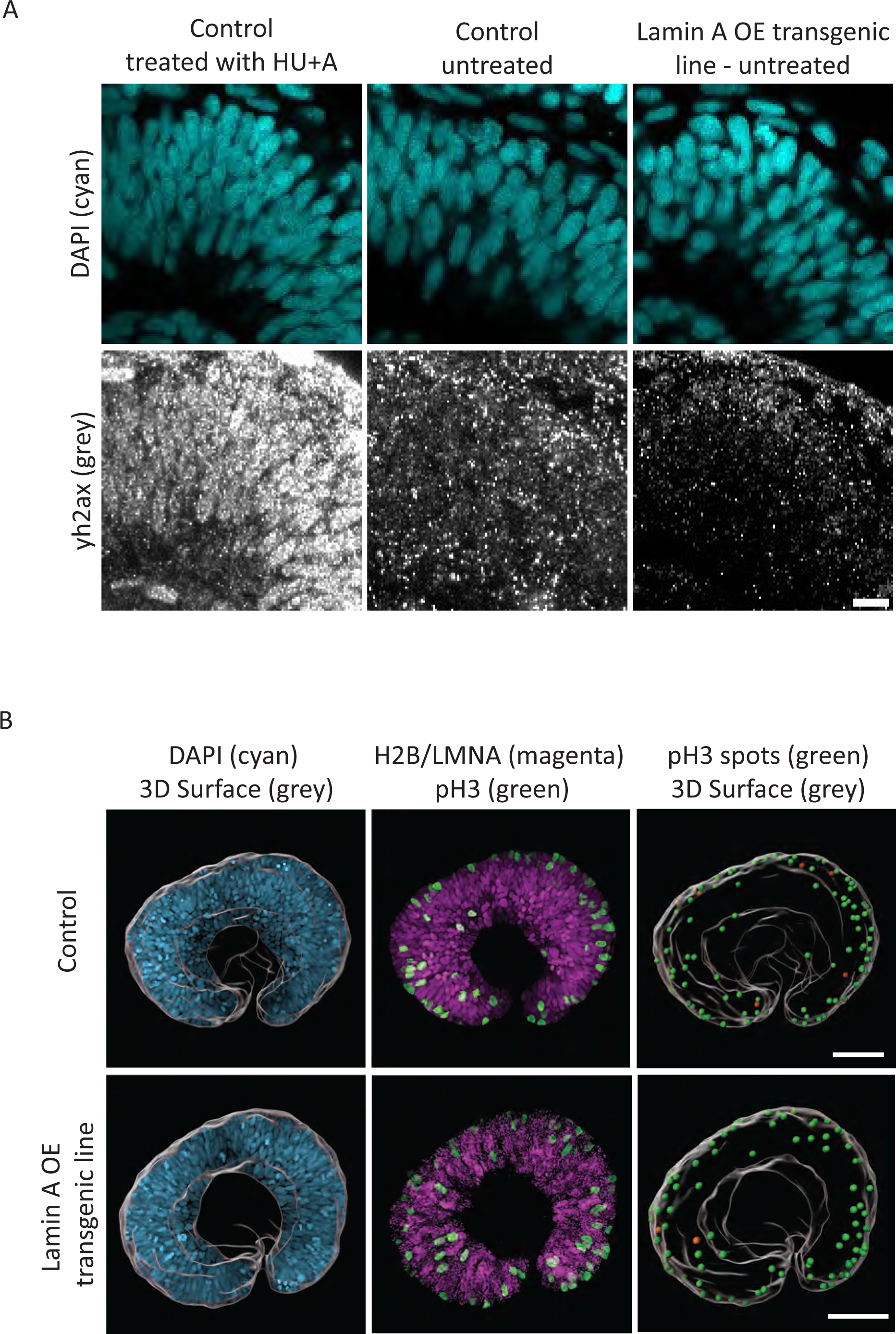
Lamin A OE nuclei show only scarce DNA damage and divide at the apical surface. (A) Immunostaining of γh2ax revealing only scarce DNA damage in control (center) and Lamin A OE transgenic line retinal neuroepithelia (right). Left: Control retinas were treated with 150 µM aphidicolin + 20 mM hydroxyurea for 6 hours to induce DNA damage (left). Scale bar: 10 μm. (B) pH3 staining (green) showing mitotic cells in the neuroepithelium of controls and Lamin A OE transgenic line (nuclei in magenta) (centre). 3D surface (grey) created from DAPI staining (left). pH3 spots (green/orange) (right) were classified according to their distance to the 3D surface. Green: pH3 spots within ≤ 5 μm from the apical surface. Orange: pH3 spots > 5 μm from the apical surface (right). Scale bar: 50 μm.

**Figure S3.**
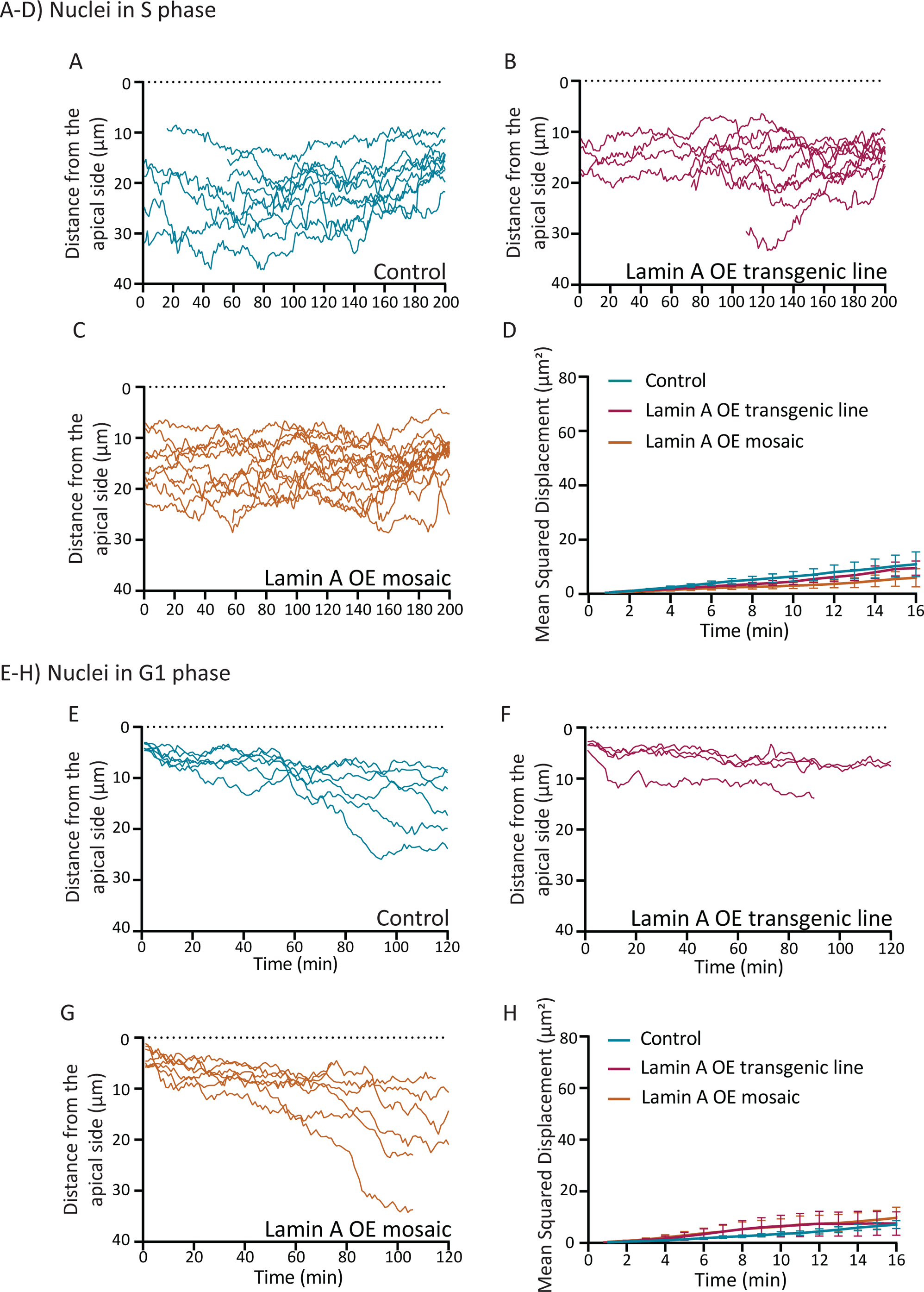
Trajectories of Lamin A OE nuclei during S and G1 phases are similar to controls. (A-C) S-phase trajectories of nuclear positions in control (A) Lamin A OE in transgenic line (B) and Lamin A OE in mosaic condition (C). t_200_ (min) corresponds to the last time-point of S phase, before G2 phase onset. (D) Mean squared displacement (MSD) of nuclear movements in S phase comparing controls (cyan) and Lamin A OE in transgenic line (magenta) and mosaic condition (orange). MSD curves mainly overlap. (E-G) G1 phase trajectories of nuclei in control (E) and Lamin A OE in transgenic line (F) and Lamin A OE in mosaic condition (G). t_1_ (min) corresponds to the time-point after cell division when daughter cells are formed. (H) Mean squared displacement (MSD) of nuclear movements in G1 phase comparing controls (cyan) and Lamin A OE in transgenic line (magenta) and mosaic condition (orange). MSD curves mainly overlap. Error bars: Standard Deviation. Imaged at 1min time intervals.

**Figure S4.**
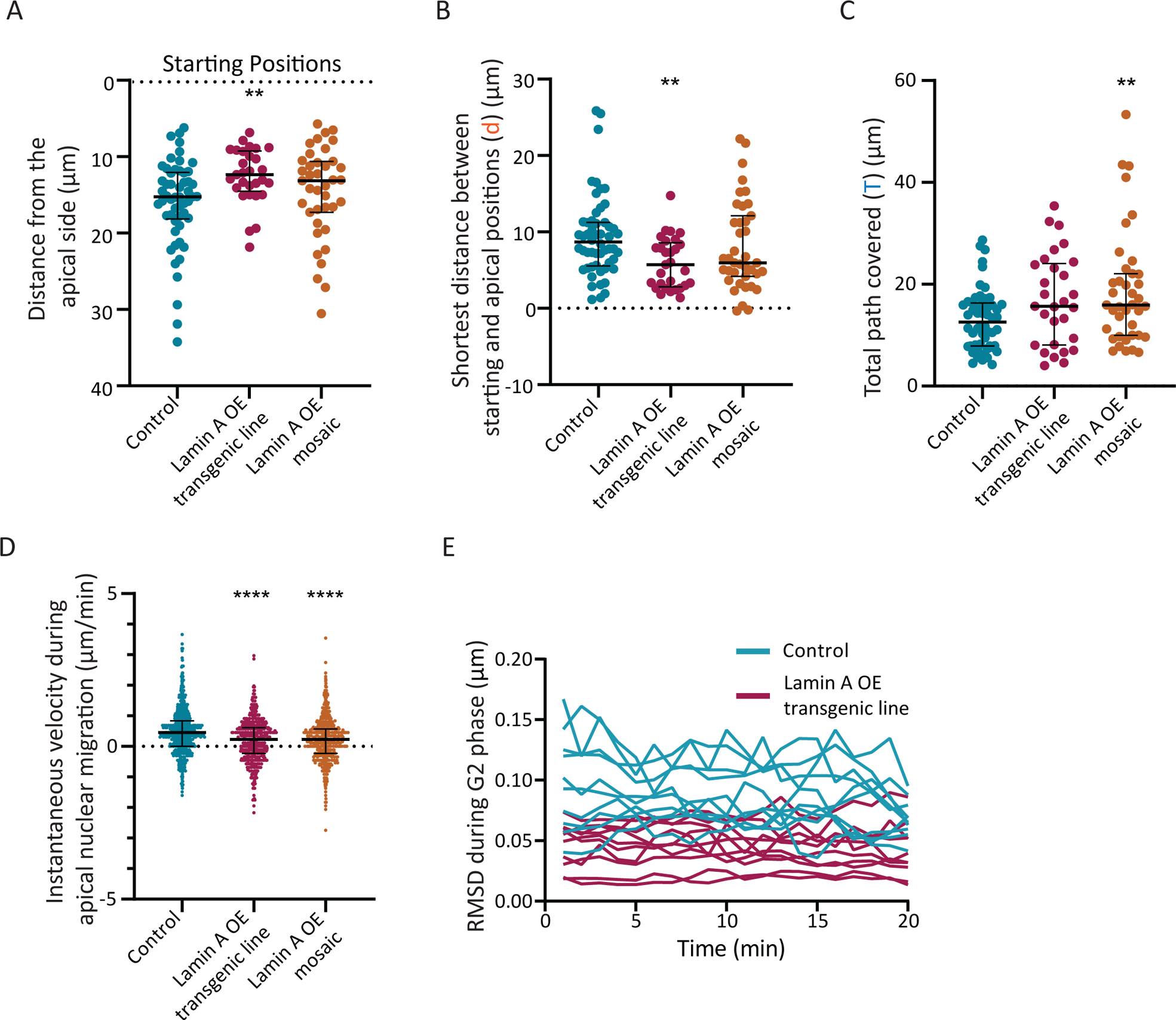
Lamin A OE nuclei are slower and cover a longer path to reach the apical surface. (A) Comparison of nuclear position at the beginning of apical nuclear migration in the control (cyan), Lamin A OE transgenic line (magenta) and Lamin A OE mosaic condition (orange). (Starting Position: P_Lamin A OE transgenic line_ = 0.0014, P_Lamin A OE mosaic_ = 0.1708, Mann-Whitney test). (B) Shortest distance between starting and division positions (d). (d: P_Lamin A OE transgenic line_ = 0.0020, P_Lamin A OE mosaic_ = 0.1566, Mann-Whitney test). Corresponds to ‘d’ in Figure 3H. (C) Total path covered during apical nuclear migration (T). (T: P_Lamin A OE transgenic line_ = 0.0720, P_Lamin A OE mosaic_ = 0.0096, Mann-Whitney test). Corresponds to ‘T’ in Figure 3H. (D) Instantaneous velocities during apical nuclear migration. (Instantaneous velocities: P_Lamin A OE transgenic line_ < 0.0001, P_Lamin A OE mosaic_ < 0.0001, Mann-Whitney test). Error bars: Median with interquartile range. (E) Control (cyan) and Lamin A OE transgenic line (magenta) trajectories of the Root Mean Squared Displacement (RMSD) for each nucleus followed during apical nuclear migration. Values were used to calculate the standard deviation in Figure 3 J.

**Figure S5.**
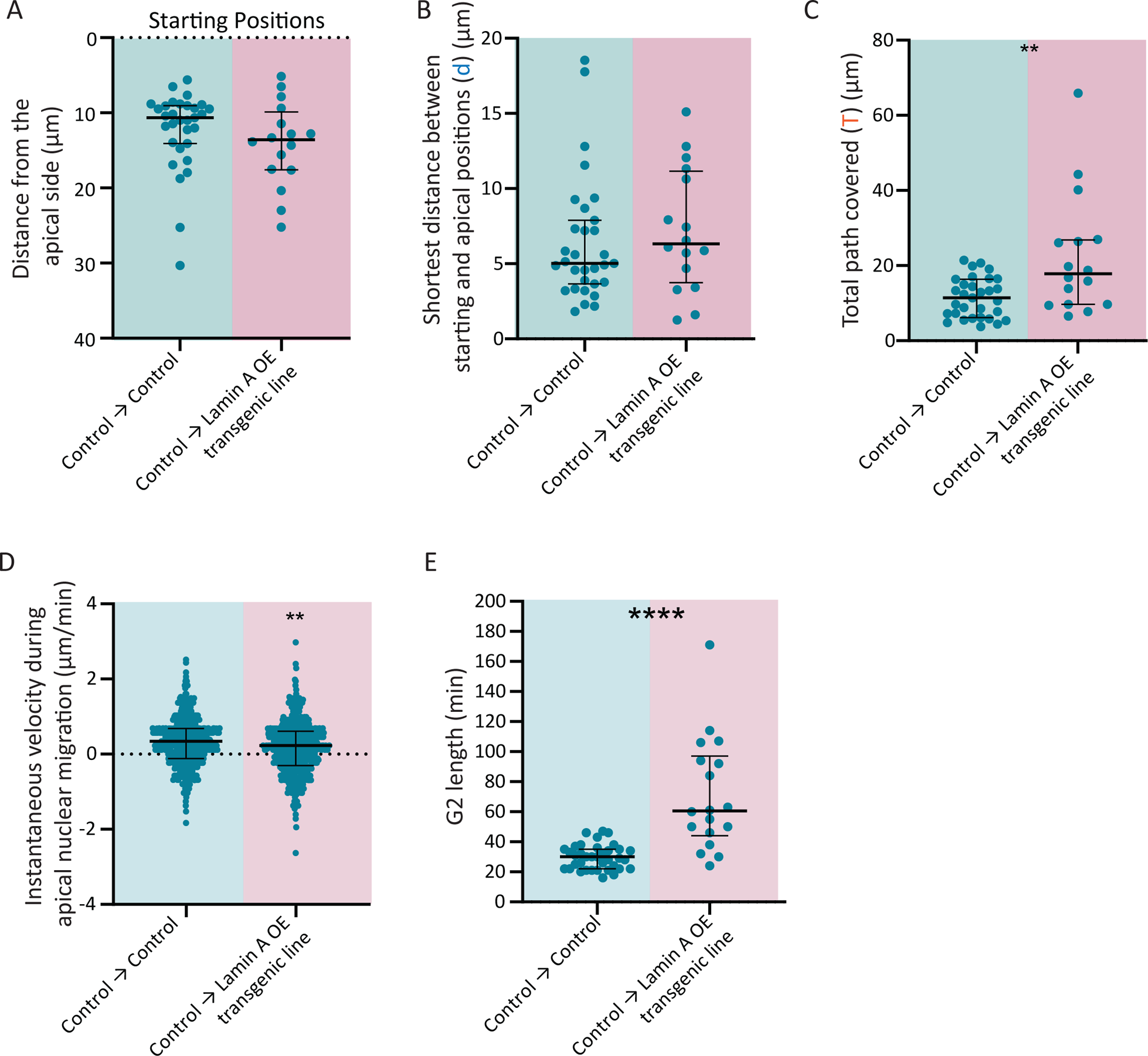
Control nuclei cover a longer path when migrating in a Lamin A OE environment. (A) Comparison of nuclear position at the beginning of apical nuclear migration when control cells are transplanted into controls or Lamin A OE line blastulas (Starting Positions: P=0,1571, Mann-Whitney test). (B) (d) Shortest distance between the starting and division positions (d: P=0,3409, Mann-Whitney test). Corresponds to ‘d’ in Figure 4G. (C) (T) Total path covered during apical nuclear migration (T: P=0,0059, Mann-Whitney test). Corresponds to ‘T’ in Figure 4G. (D) Instantaneous velocities during apical nuclear migration (Instantaneous velocities: P = 0,0015, Mann-Whitney test). (E) Length of G2 phase (min) of control nuclei transplanted into a control or a Lamin A OE environment (G2 length: P < 0.001, Mann-Whitney test). Error bars: Median with interquartile range.

## Video Legends

**Video S1. Nuclear dynamics in the densely packed zebrafish retina.**

Example of nuclear movements in the crowded zebrafish retinal neuroepithelium. Imaging of the Tg(hsp70:H2B-RFP) zebrafish line which labels nuclei (grey). Time interval = 1min. Scale bar = 10 μm.

**Video S2. Apical nuclear migration in the control, Lamin A OE transgenic line and Lamin A mosaic conditions.**

Example of apical nuclear migration in control, Lamin A OE transgenic line and Lamin A OE mosaic conditions. Nuclei are labelled with PCNA-GFP (cyan). Nuclei overexpressing Lamin A are labeled in magenta. Red dot indicates the nucleus followed. Time interval = 1min. Scale bar = 10 μm.

**Video S3. Apical nuclear migration of control nuclei into a control and Lamin A OE environment.**

Example of control nucleus (grey) during apical nuclear migration in a Control (cyan) or Lamin A OE transgenic (magenta) environment. Transplanted nuclei are labelled with PCNA-GFP (grey). Control nuclei are labeled in cyan and nuclei overexpressing Lamin A are labeled in magenta. Red dot indicates the nucleus followed. Time interval = 1min. Scale bar = 10 μm.

**Video S4. Nuclear envelop breakdown of control nuclei into a control and Lamin A OE environment.**

Example of control nucleus (grey) reaching the apical surface to nuclear envelop breakdown in a Control (cyan) or Lamin A OE transgenic (magenta) environment. Transplanted nucleus is labelled with PCNA-GFP (grey). Nuclei of recipient cells are labelled with hsp70:H2B-RFP (cyan) or with hsp70:LMNA-mKate2 (magenta). Red dot indicates the nucleus followed. Time interval = 1 min. Scale bar = 10 μm.

**Video S5. Mitotic rounding of control cells into a control and Lamin A OE environment.**

Example of control cells reaching the apical surface in a Control or Lamin A OE transgenic environment. Plasma membrane is labelled with RAS-GFP (grey). Control → Control: transplanted cells show the nucleus in cyan and the plasma membrane in grey; cells of recipient show the nucleus in cyan. Control → Lamin A OE transgenic line: transplanted cells show the nucleus in cyan and the plasma membrane in grey; Recipient cells overexpressing Lamin A are not visible. Red dot indicates the nucleus followed. Time interval between frames = 1min. Scale bar = 10 μm.

**Video S6. Nuclear Laser Ablation**

Example of fluid-like behavior of nuclei upon laser ablation in the tissue (by Iskra Yanakieva^35^). Nuclei are labelled with H2B-RFP. Time interval = 10 seconds.

## SUPPLEMENTARY NOTE

### 1 Descriptors of nuclear shapes in 3D

The 3D shapes of nuclei were approximated by a surface triangulation (see Nuclear segmentation, volume and shape in Materials and Methods). We fitted the equation of an ellipsoid

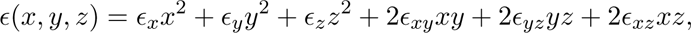

to the geometric centers ***p****_i_*_=1_,_2_*_…N_* of *N* triangular simplices on the nuclei’s surfaces in the Cartesian barycentric coordinates 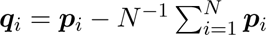 by minimizing the least-squares error

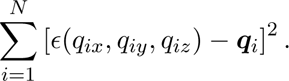

In the left-handed eigensystem associated with the inverse matrix *M^−^*^1^, which is determined by

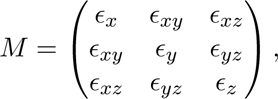

the fitted ellipsoid is described by three elliptical axes *e_x_*, *e_y_*, and *e_z_* (the square roots of the eigenvalues of *M^−^*^1^) with the major axis aligned along *z*.

To characterize the average deviation of the nuclear shapes from the perfect ellipsoid, we transformed the Cartesian coordinates (*x, y, z*) associated with the eigensystem of *M^−^*^1^ into the spherical coordinates (*r, θ, ϕ*), in which the best-fit ellipsoid is given by (*ɛ, θ, ϕ*) with

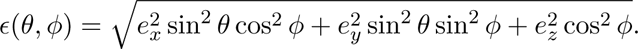

Then we evaluated the root mean squared deviation as

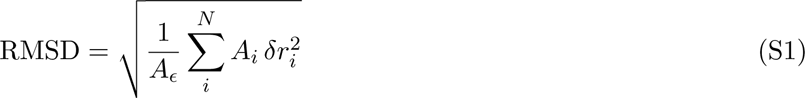

with *δr_i_* = *r_i_ ɛ*(*θ_i_, ϕ_i_*), in which *A_ɛ_*, *A_i_*, and (*r_i_, θ_i_, ϕ_i_*) are, respectively, the total area of the ellipsoid *ɛ*(*θ, ϕ*), the area of the *i*^th^ triangular simplex, and the spherical coordinates of its geometric center ***q****_i_*. The formula Eq. S1 for the RMSD represents a numerical approximation for the integral

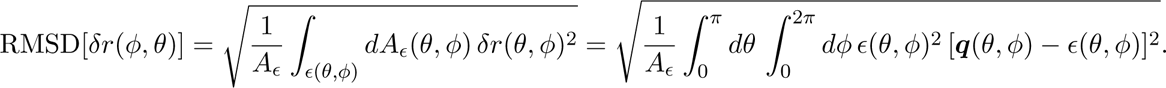

The rationale behind the definition of RMSD is to provide an average measure of deviations from a reference shape, which does not depend on the total area (size) of the latter. For example, consider that we have an ellipsoid *ɛ*. Then we perturb its shape by a constant deviation *δɛ*. The RMSD calculated for the reference shape *ɛ* then yields

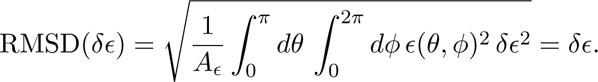

### 2 Model of a compressible droplet for nuclear shape

Our system of interest is nearly axisymmetric with nuclei stretched along the cell axis perpendicular to the epithelial plane. We therefore assume that nuclear shapes can be approximated as prolate spheroids with longer axis *a* = *e_z_* and shorter axis 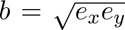. The geometric mean of the minor axes ensures that the spheroid’s volume

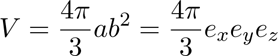

equals the volume of the best-fit ellipsoid characterized by the axes *e_x_*, *e_y_*, and *e_z_* (Sec. 1).

We extend a common soft-matter model for the free energy of a compressible droplet [3, 4, 6, 2, 1]

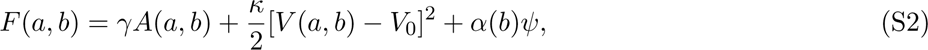

in which *A*(*a, b*) is the spheroid surface area, and the constants *κ* and *V*_0_ are, respectively, volume elasticity and preferred volume of the droplet. Additionally, we introduce a term that penalizes the spheroid’s cross-sectional area to account for compressive effects in the plane of the surrounding tissue, in which the coefficient *ψ* is conjugate to the cross-sectional area *α*(*b*) = *π*^2^*b*^2^. Because there is no closed-form expression for *A*(*a, b*) which is valid for both prolate and oblate spheroids, we approximate the surface area by a generalized mean

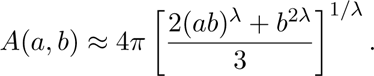

This class of approximations, *P* (*λ, λ,* 0) in the notation of Ref. [5], is exact for spheres and leads to a very small relative error when *λ ≈* 8*/*5—a value attributed to an unpublished result of Knud Thomsen.

We extract the parameters of the model from the time series of the axes *a* and *b*, which were obtained by fitting the triangulated surfaces of nuclei (Sec. 1, Nuclear segmentation, volume and shape in Materials and methods). We assume that S-phase nuclei fluctuate about their equilibrium shapes, which are characterized by

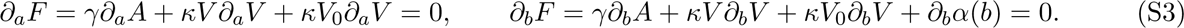

We normalize the above equations by the compression coefficient *ψ >* 0, which stabilizes the prolate spheroid in equilibrium, so that we can estimate the mechanical parameters relative to the compressive tension. We optimize *γ/ψ*, *κ_ψ_* = *κ/ψ* and *κ*_0_ = *κV*_0_*/ψ* over observations *{*(*a, b*)*_i_*_=1_,_2_,.} to fit Eq. (S3) in the least-squares sense for each nucleus individually. The value of the preferred volume is inferred from *V*_0_ = *κ*_0_*/κ_ψ_* afterwards.

